# Representations of Pitch and Timbre of Instrument Sounds in the Inferior Colliculus

**DOI:** 10.64898/2026.08.09.743816

**Authors:** Johanna B. Fritzinger, Laurel H. Carney

**Author notes:** Corresponding Author: Johanna B. Fritzinger.

## Abstract

**Purpose:** The neural representation of pitch and timbre in complex sounds has previously been studied using synthetic, controlled stimuli to investigate underlying encoding mechanisms. These studies provide information about how single attributes of sound are represented in the inferior colliculus (IC), a critical hub of the auditory pathway where neurons are sensitive to stimulus periodicity and spectral shape, giving rise to representations of pitch and timbre, respectively. However, there is a gap in understanding how natural sounds with both pitch and timbre attributes, such as instrument sounds, are represented in the IC.

**Methods:** In this study, extracellular recordings were made in the IC of awake rabbits in response to natural instrument stimuli varying in fundamental frequency (F0) to determine how instrument identity (timbre) and F0 (pitch) are represented in IC neurons.

**Results:** Using decoding models for instrument identification, we found that instrument identity was redundantly encoded in a population of neurons with diverse rate and timing characteristics. F0 identification using decoding models trained on single-neuron rate responses was poor, but the population of rate responses contained enough information to identify F0 reliably. F0 information was also encoded in single-neuron temporal responses up to 196 Hz. F0 identification from a population of temporal responses was accurate up to approximately 900 Hz, but accuracy decreased at high F0s. For the task in which F0 was identified based on responses to both oboe and bassoon stimuli that had overlapping F0s, performance decreased compared to F0 identification based on responses to a single instrument.

**Conclusion:** This result supports the hypothesis that pitch and timbre information are encoded jointly in the IC.

## Introduction

Pitch and timbre are critical aspects of speech and music. Previous studies used synthetic sounds with flat or triangular-shaped spectral envelopes to explore how neurons in the inferior colliculus (IC) encode these acoustic features. However, how IC neurons respond to natural instrument sounds, which have more spectral peaks and more complex phase spectra than the synthetic stimuli used in previous studies, is unknown. This study aimed to fill this gap by recording IC responses in awake rabbit to bassoon and oboe stimuli with varying fundamental frequencies (F0). Bassoon and oboe were chosen because their spectral envelopes are simpler than many other instruments and more similar to the triangular envelope used in previous synthetic studies [1]. This study investigated how neurons in the IC encode F0, instrument identity, and their interaction.

Studies using synthetic sounds to investigate one characteristic, F0 or timbre, provide the basis for initial hypotheses about natural instrument encoding. F0 encoding has been studied in the IC of awake rabbits using flat-spectrum, harmonic-tone complexes (HTCs) [2, 3]. Behaviorally, rabbits discriminate F0 using temporal envelope cues at low F0s and spectral cues of resolved harmonics at high F0s [4]. Analysis of IC responses to these stimuli supports encoding of F0 based on average rate (for F0s > 800 Hz), phase-locking to envelope (F0s < 900 Hz), and, for neurons excited by amplitude modulations, by band-pass rate tuning to envelope repetition rate (200 – 1600 Hz) [2, 3].

Timbre- and vowel-encoding studies have also used synthetic stimuli to investigate how spectral peaks are represented in the IC [1]. HTCs with broad triangular spectral peaks and zero-phase spectra are encoded in the average discharge rates of IC neurons [1]. Similarly, formant discrimination based on average-rate information in budgerigar IC is sufficient to describe behavioral formant discrimination in quiet, though timing information was necessary when background noise was added [5]. Timing information was also necessary for single-neuron identification of spoken vowels, and vowel accuracy was correlated with neural sensitivity to low-velocity chirps [6]. Another study found that IC models with amplitude-modulation sensitivity accurately predicted some neural responses to spoken vowels [7]. These studies were limited to F0s near the voice-pitch range; higher F0s, which are important to consider in studies of natural instrument sounds, may be encoded by different mechanisms.

By using natural instrument sounds with a range of F0s, this study was also designed to explore an open question about pitch and timbre interactions. Psychophysical studies using synthetic stimuli [8] and natural vowel and instrument stimuli [9] have shown that pitch impacts timbre discrimination performance, and vice versa. Modeling work has proposed that this interaction may be present in the IC [10]. The spectral centroid, a measure of timbral brightness, varies with F0 within a single instrument [11], thus complicating pitch and timbre separability in natural sounds.

In this study, we hypothesized that rate information across a population of IC neurons is sufficient to classify instrument identity, based on previous timbre-discrimination results in quiet [1]. For F0 identification, we hypothesized that timing information is necessary for F0s < 900 Hz, but that rate-based decoders would be accurate at high F0s, in part due to changes in spectral centroid for different F0s. We hypothesized that neurons with sensitivity to amplitude modulation could encode both F0 and instrument identity. Previous cortical studies have shown that pitch and timbre information of vowels are often represented in the same neurons [12]. It is unknown whether this interaction occurs in the IC, and our study aimed to investigate that question as well.

Single-neuron responses were recorded in the central nucleus of the IC in awake rabbits to bassoon and oboe sounds with F0s ranging from 58–1661 Hz. First, the variability in these stimuli was analyzed. Then we investigated whether a decoding model trained on a single neuron, or a population of neurons, could accurately identify instrument and/or F0 using rate or timing information. We used classification and linear regression models to study information present in the IC, determine if specific IC neuron responses are necessary for identification, and investigate overlap in neurons that contain information about pitch and instrument identity. We found that a population of neurons with diverse characteristics was sufficient to encode instrument identity, and both rate and timing information in populations of IC units could encode F0. Often, single-neuron responses that contained information about lower F0s also contained information about instrument identity.

## Methods

### Animal Care and Procedures

Extracellular, single-unit recordings were made in the central nucleus of the IC in four female Dutch-belted rabbits from 5 months to 5 years of age. All methods were approved by the University of Rochester Committee on Animal Resources. Surgical procedures consisted of an initial headbar placement, craniotomy and microdrive placement, and subsequent microdrive replacements. Animals were anesthetized intramuscularly with 66 mg/kg ketamine and 2 mg/kg xylazine, or 35mg/kg ketamine and 0.10-0.15mg/kg dexmedetomidine for all procedures. An initial surgery was performed to affix a custom, plastic, 3D-printed headbar (ProtoLabs, Maple Plain, MN) to the skull using screws and dental acrylic. One month after the initial surgery, another surgery was performed to create the craniotomy and insert the microdrive into the headbar. Subsequent surgeries were performed every 3-6 months to replace the microdrive and change position to sample different areas of the IC. Normal hearing was monitored using distortion-product otoacoustic emissions (Whitehead et al., 1992). When emissions dropped by 10 dB, experiments were discontinued, and animals were perfused to extract the brain. The IC tissue was stained with Nissl to confirm tetrode location in the IC.

### Physiological Recordings

Rabbits were trained to sit in a custom chair with their heads fixed using the headbar. They were placed inside a sound-attenuated booth (Acoustic Systems, Austin, Texas, USA). Daily recording sessions lasted up to two hours.

Calibration tones and stimuli were created in MATLAB and sent to an audio interface (16A, Mark of the Unicorn, Cambridge, Massachusetts, USA). The stimuli were converted from digital to analog (DAC3 HGC, Benchmark Media Systems, Inc., Syracuse, New York, USA) and presented through a speaker (Beyerdynamic DT-48, Beyerdynamic GmbH and Co., Heilbronn, Germany or Etymotic ER2, Etymotic Research, Inc., Elk Grove Village, Illinois).

At the beginning of each session, custom earmolds (Hal-Hen Company, Inc., Garden City Park, NY, USA or Dreve Otoform Ak, Unna, Germany) were inserted and the frequency response of the acoustic system was calibrated. Tones of 50–20,000 Hz were presented and a probe-tube microphone (Etymotic ER10B+ or ER7C, Etymotic Research, Inc., Elk Grove Village, Illinois) was used to record in the ear canal. All stimuli were filtered to compensate for the frequency response of the system.

Neural signals were recorded using a microdrive that contained four tetrodes. Tetrodes consisted of four twisted 18-μm epoxy-coated platinum-iridium wires (California Fine Wire Co., Grover Beach, CA) plated with a platinum-black solution (Neuralynx, Inc., Bozeman, MT) to lower impedances to approximately 0.1-1.5 MOhms. The tetrodes were advanced or retracted at the end of the sessions using the microdrive to sample different areas of the IC. Signals from the tetrodes were amplified (RHD2216 16-channel amplifier chip) and recorded using the Intan RHD recording system (Intan Technologies, LLC., Los Angeles, CA, USA) and accompanying software. Signals were recorded at 30-kHz sampling rate and filtered with a 4th-order Butterworth 300 Hz-3 kHz filter.

After each recording session, signals were sorted using custom MATLAB code to identify single neurons. The thresholds for each wire were set at 4 times the standard deviation of the waveform [13]. Waveforms were sorted into individual neurons using custom software [14]. Waveforms had to meet 2 criteria to be considered single-neuron recordings: 1) the percent of spikes that occurred with intervals less than 1 ms was less than 2%, and a cluster entanglement metric had to be less than 0.1 [14]. Spike times were extracted and saved.

### Stimuli & Analysis

Several basic stimuli were included to characterize each neuron. First, pure tones from 250–16,000 Hz were presented in 5 steps per octave to classify the CF and response map of each neuron. Tones were presented at 10, 30, 50 and 70 dB SPL with a 200-ms duration and 10-ms raised-cosine on/off ramps. Stimuli were presented diotically in random order for three repetitions with 400-ms interstimulus intervals. The CF of the neuron was defined as the frequency that elicited an increase in average rate at the lowest sound level.

Modulation transfer functions (MTFs) in response to amplitude-modulated (AM) stimuli were characterized by presenting sinusoidally amplitude-modulated gaussian noise. The noise bandwidth was 100 Hz to 10 kHz, modulation frequencies spanned 2–600 Hz in 3 steps per octave. AM stimuli were presented at 33-dB SPL spectrum level, 1-s duration with 50-ms raised-cosine on/off ramps, and 500-ms interstimulus intervals. The stimuli were repeated 5 times and were presented diotically in a random order. Average-rate responses were calculated, excluding a 50-ms onset response. Neurons were classified into 5 types based on the comparison of responses to modulated and unmodulated noises: band-enhanced (BE), band-suppressed (BS), hybrid (BE-BS), hybrid (BS-BE), and flat. BE neurons had at least 2 modulation-frequency responses with rates significantly higher than the unmodulated rate (two-sample t-test, p<0.05), without a response at an intermediate modulation frequency that was significantly below the unmodulated rate. BS neurons had at least 2 modulation-frequency response rates significantly below the unmodulated rate, again without a significantly higher response rate at an intermediate modulation frequency. Hybrid neurons met both BE and BS criteria. H (BE-BS) neurons were BE at low modulation frequencies and BS at higher modulation frequencies, whereas H (BS-BE) were the opposite. Flat neurons did not meet either BE or BS criteria. The best modulation frequency (BMF) and worse modulation frequency (WMF) was calculated from the MTFs of BE and BS neurons, respectively. The BMF/WMF was the modulation frequency with the maximum/minimum of the spline-interpolated average rate response.

Neural sensitivity to fast frequency sweeps, known as chirps, was characterized by rate-velocity functions (RVFs) [6, 15] based on stimuli that consisted of chirps with velocities of 0.25, 0.5, 0.75, 1, 1.25, 1.5, 1.75, 2, 2.5, 3, 4, 5, 6, 7, 8, 9 kHz/ms, in both the positive and negative directions. Stimuli were normalized by energy and included raised-cosine on/off ramps that were 10% of the duration of each chirp. Chirp duration varied based on velocity. Chirps were presented in random order, diotically, with a random interstimulus interval of 40-60 ms. Each chirp was presented 80 times. Response rates were based on sums over a 15-ms time window that started at an estimate of neural latency, defined as the latency for tones at CF at 73 dB SPL. Principal components analysis was used to extract prominent features of the chirp responses, with the first three principal components matching previous work [15].

The natural stimuli consisted of samples of musical instrument sounds (University of Iowa Electronic Music Studios, Iowa City, IA). A 300-ms duration, steady-state segment of the sound was extracted, and 20-ms raised-cosine ramps were added to the beginning and end of the stimulus. Stimuli were presented diotically at 73 dB SPL, in random order, for 20 repetitions. The bassoon stimulus set included all semitones from Bb1 (58 Hz) to D5 (587 Hz), and the oboe stimulus set included all semitones from Bb3 (233 Hz) to Ab6 (1661 Hz). Sixteen of the bassoon stimuli had F0s that overlapped with the oboe.

The spectral centroid of each stimulus was calculated to quantify changes over F0. To calculate the spectral centroid of a stimulus, the fast-Fourier transform was computed with a resolution of 3.33 Hz. The spectral centroid in Hz was calculated as:

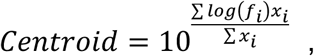

where *x_i_* is the log amplitude of the frequency spectrum at bin i, and *f_i_* is the logarithmically transformed frequency of that bin. The F0 values of the stimuli were also calculated from the stimulus spectrum, as the labeled F0 from the recordings were not always accurate. F0 was estimated first by finding the frequencies of the stimulus harmonics using Welch’s power spectral density estimate and finding the peaks of the spectrum. Then, an initial estimate of F0 was defined as the first spectral peak. The estimate was then refined iteratively by taking the difference between the next harmonic and previous harmonic and averaging that difference with the current F0 estimate. This process continued for harmonics up to 10 kHz.

Neural responses to natural-timbre stimuli were analyzed by calculating rate and timing metrics. Average-rate responses were computed over the duration of the stimulus, averaged over all repetitions, excluding a 50-ms onset response. Additionally, classification analyses required calculating average rate per repetition of the stimulus. Temporal analyses included calculating peri-stimulus time histograms (PSTHs), period histograms, inter-spike intervals, vector strength, and reliability. Vector strength (VS) was calculated as:

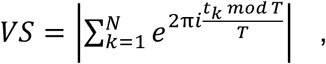

where *t_k_* are spike times for all repetitions, excluding the 50-ms onset, and T is the period of the harmonic of interest. The time of each spike within a period was calculated based on the spike time modulo the period. Vector strengths to F0 and to individual harmonics were calculated. Significance of vector strength was determined using a standard Rayleigh test (p<0.01).

A reliability metric was used to determine consistency of spiking activity of a neuron over many repetitions [16]. Reliability was defined as the correlation coefficient between the PSTHs of even and odd repetitions of the stimulus.

### Classification Tasks

Three different decoding tasks were performed using various classification analyses and decoding models. The first task was identifying instrument identity, the second task was identifying F0 of either oboe or bassoon, and the third task involved identifying both instrument identity and F0 of the stimulus. The instrument-identification task included both oboe and bassoon stimuli that overlapped in F0, which included 16 stimuli with F0s ranging from 233–587 Hz. The goal of the analysis was to correctly identify the instrument identity regardless of F0. The F0-identification task was split into identifying F0 based on responses to the bassoon stimuli (40 stimuli), identifying F0 based on responses to the oboe stimuli (35 stimuli), or identifying F0 in the overlapping F0 range using responses to both bassoon and oboe (16 stimuli). Two other F0 identification tasks were performed: 1) training on a subset of F0s and predicting the held-out F0s, and 2) training on 16 overlapped oboe stimuli and testing on the 16 bassoon stimuli, and vice versa. Lastly, the instrument-identity-and-F0-discrimination task used responses to all 75 stimuli, and the goal was to determine if neurons or populations of neurons could correctly identify both instrument and F0. All tasks used the same types of analysis and models, presented below.

### Single-Neuron Classification Analyses

First, a simple rate metric was used to determine how well a single-neuron rate response could classify either instrument identity and/or F0 using a leave-one-out procedure (Mitchell and Carney, 2025). For the F0 identification, the average-rate response for a single repetition of one F0 stimulus was calculated. Then, the average-rate response across the other 19 repetitions of the stimulus and across the 20 repetitions of all other F0 stimuli provided an array of overall-average rates for all F0 stimuli. The single repetition response was compared to the overall average rates. The F0 classification of the single average rate response was determined as the stimulus with an overall-average rate closest to the single-repetition rate. This analysis was performed for all repetitions and all conditions and for all neurons. Instrument identity and F0/instrument identity rate discriminations were performed in the same manner. A confusion matrix was calculated based on the classifications. Overall accuracy was defined as the sum of the diagonal of the confusion matrix (correct responses) divided by the total number of responses. Accuracy was calculated for all classification tasks.

Next, classification models were used to test how well timing information from a neuron could classify instrument identity and/or F0. Model features included binned spike times from each trial, where the bin width was varied from 0.1 ms to 300 ms (the average-rate response) and the bin width was chosen as the value that resulted in the highest accuracy. All models were trained on 80% of the data and tested on the other 20%. For these tasks, the training data was chosen as 16 random repetitions out of the 20 stimulus repetitions to ensure stimulus classes were equally represented in the training data. Support vector machines (SVM) were used as the base learning algorithm. Five-fold cross validation was performed on the training data to provide robust estimates of model performance. The regularization strength, lambda, was chosen by systematically varying lambda and using the lambda value that resulted in the lowest mean squared error in the final model. Ridge regularization was used to prevent overfitting and improve model generalization. The convergence criterion was set to 0.0001 for stable model convergence. The instrument identification task used the MATLAB function fitclinear, the F0 task used fitcecoc.

### Population Classification Analyses

Population analyses included decoders that used the average-rate responses or PSTH information to test rate vs timing predictions in a population of IC neurons. For all tasks, the models were trained on 80% of the data and tested on the remaining 20%, again stratified such that 16 reps of each stimulus were used in training. All decoders used the MATLAB function fitcecoc. The learning algorithm was SVM, using a multi-class strategy of error-correcting output codes to predict the classifications. Ridge regularization was used to prevent overfitting and improve generalization, and the optimal regularization strength (lambda) was optimized as mentioned previously. Five-fold cross validation was performed on the training data. An analysis of feature weights, called beta weights, was performed to investigate the contribution of each neuron to the model. Timing information in a population of neurons was used for the F0-decoding task and F0- and-instrument-identification task, using PSTHs from 1–40 neurons with a bin size of 1 ms to keep computation time down. This model was not used for the instrument-identification task because discharge rates were sufficient to accurately predict instrument. Permutation testing was performed to determine the importance of each neuron in the model.

### Regression Analyses for the F0-Identification Task

Lastly, linear regression models were tested for the F0-identification task. Regression models were fit to the single-neuron PSTH data (bin size of 0.25 ms), the population-rate data, and the population-PSTH data (bin size of 1 ms). All models were fit using the MATLAB function fitrlinear. An 80/20 train/test split was used for model evaluation. For the basic F0-estimation tasks, the training/testing sets were stratified such that for each set of 20 repetitions, 16 (80%) repetitions were used in training and 4 (20%) were used in testing. Least-squares regression was selected as the learning algorithm. Ridge regression (L2) was applied to prevent the models from overfitting. The regularization strength, lambda, was chosen by systematically varying lambda and using the lambda value that resulted in the lowest mean squared error in the final model. The limited-memory Broyden-Fletcher-Goldfarb-Shanno solver was used for parameter estimation. A convergence criterion was set to 0.001 for the beta coefficients to maintain efficiency. Two other training/testing configurations were performed. The first configuration trained the linear regression models on 80% of the F0s and tested the model using the remaining 20% of the F0s. The F0s used for training/testing were randomly selected. The second task trained on the 16 overlapping F0s for one instrument (50% training data) and tested on the 16 overlapping F0s for the other instrument (50% testing data). The model was fit using the regression methods described above.

### IC Computational Models

Three computational models of IC neurons were evaluated to determine their ability to predict the neural responses. Detailed implementations of all three models can be found in [17]. The first model was an energy model consisting of a 4^th^-order gammatone filterbank with cat Q10 tuning values. The other two models used an auditory-nerve model stage [18]. The second model was the same-frequency excitation-inhibition (SFIE) model, which consists of a VCN stage and IC BE and IC BS cell stage that accurately predict IC sensitivity to amplitude modulation [19, 20]. The last model was a modified SFIE model that included additional inhibition from two off-CF BS cells [17] to simulate broadband modulation-sensitive inhibition (BMSI). The BMSI model was developed to predict BE and BS MTF types, as well as IC responses to tones in wideband noise [17] and synthetic timbre stimuli [1].

For the energy, SFIE, and BMSI models, model predictions were made for each recorded neuron, with parameters selected based on CF and MTF type. Model average-rate responses were compared to data using the squared correlation coefficient (R^2^) between model and observed responses. The amount of neural variance explained by the model rates was above chance if the R^2^ value was significantly above an R^2^ value based on model rates that had been shuffled. For the SFIE model, a non-homogenous Poisson process was used as a spiking generator, and spiking activity was calculated for each model neuron. SFIE spiking activity was used as the input to the F0-discrimination and instrument-identification decoding models to analyze differences between performance based on model responses and IC responses.

### Statistics

Statistical tests included linear regression to determine relationships between model accuracies or model weights and features such as CF, second principal component of the RVF, phase-locking strengths, and spiking reliability. Kruskal-Wallis for non-parametric data and Mann-Whitney U-tests were also used to determine significant differences in accuracies due to PSTH bins or MTF groups. Variance explained, R^2^, was used in linear regression models to compare model performance to discriminations or identifications based on the neural data.

### Code Accessibility & Data Availability

Custom MATLAB code for clustering data (Schwarz et al., 2012) is available at https://www.urmc.rochester.edu/labs/carney/publications-code/spike-sorting-code.aspx. Data and custom MATLAB code for data analysis and modeling will be available on OSF. Code is also available on GitHub at https://github.com/jfritzinger/FritzingerCarney2026-NatTimbre.

## Results

For the stimulus set used here, the bassoon and oboe stimuli had different spectral-peak locations, and the oboe had more energy in the higher harmonics (Fig. 1a) for all overlapping F0s. For both oboe and bassoon, spectral centroid increased as a function of F0 (Fig. 1b). Oboe stimuli also had a higher spectral centroids compared to bassoon, eliciting a brighter timbre (Fig. 1b), as expected.

**Figure 1.**
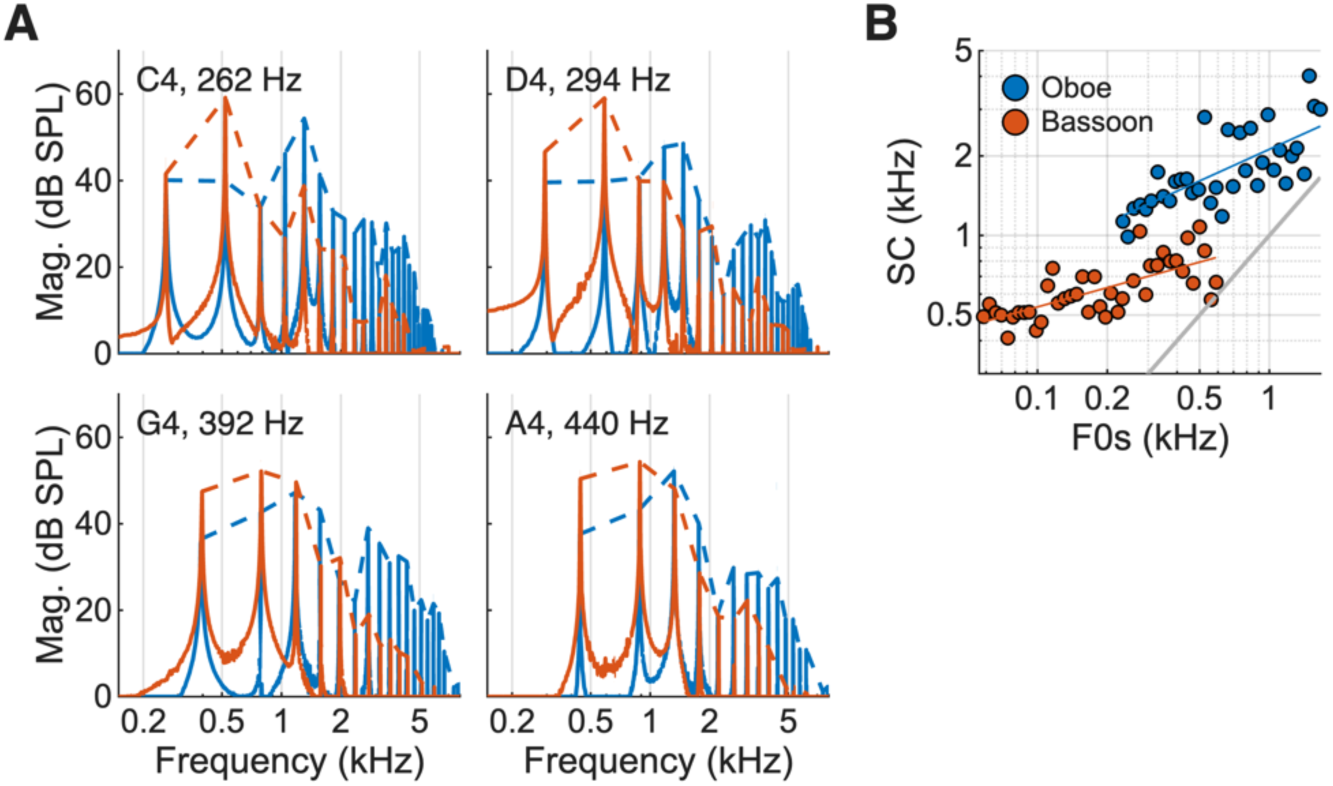
Characteristics of the natural-timbre stimuli. (A) Four examples of oboe and bassoon magnitude spectra for stimuli with different F0s. (B) Spectral centroid (SC) plotted as a function of F0 for oboe and bassoon stimuli.

Single-neuron responses were recorded from a total of 298 neurons in the central nucleus of the IC. Responses to both bassoon and oboe stimuli were collected for 249 neurons. Bassoon responses only were recorded for an addition 41 neurons and oboe responses only were recorded for an additional 7 neurons. Response maps and MTFs were collected for all neurons, and RVFs were collected for 188 neurons. Neural CFs ranged from 332–13,900 Hz (Fig. 2a). MTFs were analyzed for the population of responses: 22% (n=67) were BE, 49% (n=146) were BS, 18% (n=54) were hybrid, and 11% (n=31) were flat (Fig. 2b). The median BMF for BE neurons was 83 Hz, and the median WMF for BS neurons was 76 Hz (Fig. 2c, d).

**Figure 2.**
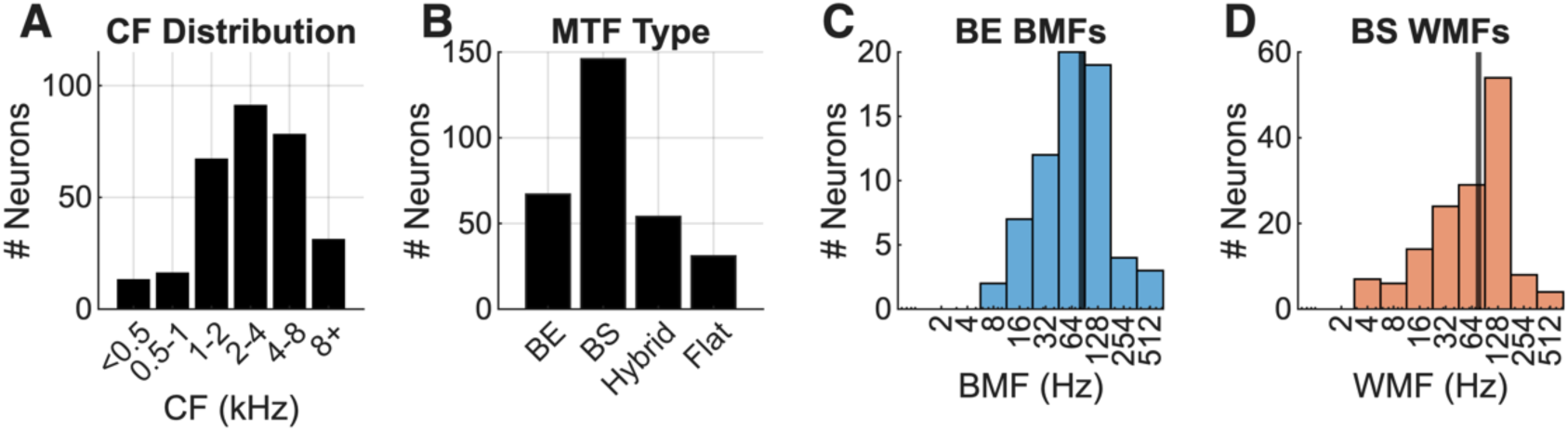
Distributions of neural metrics (n = 298 neurons). (A) CFs; (B) MTF types; (C) Best modulation frequencies (BMFs) for BE neurons, n=67; (D) Worst modulation frequencies (WMFs) for BS neurons, n=146.

### Instrument Identification

#### Identification of instrument based on individual-neuron rate and timing information

The first task was decoding instrument identity over the range of overlapping F0s, 233–587 Hz, using rate or timing information from single neurons. Instrument-identification accuracy based on rate responses from single neurons had a mean accuracy of 59%, ranging from 34 to 88%. The identification accuracy of 16 neurons was below chance (50%) reflecting an unreliable classification boundary driven by high variance in the data (Fig. 3d). Rate responses from 12 example neurons revealed that neurons with the highest accuracy had consistent rate differences over the range of overlapping F0s (Fig. 3a, b, c); in contrast, neurons exhibiting chance accuracy showed overlapping response rates (Fig. 3d). Identification accuracy was not correlated with CF and did not differ significantly across MTF types (Fig. 3e). Because differences in CF and MTF type could not account for identification accuracy, we next tested whether stimulus energy differences between instruments were driving neural rate differences (Fig. 3f)

**Figure 3.**
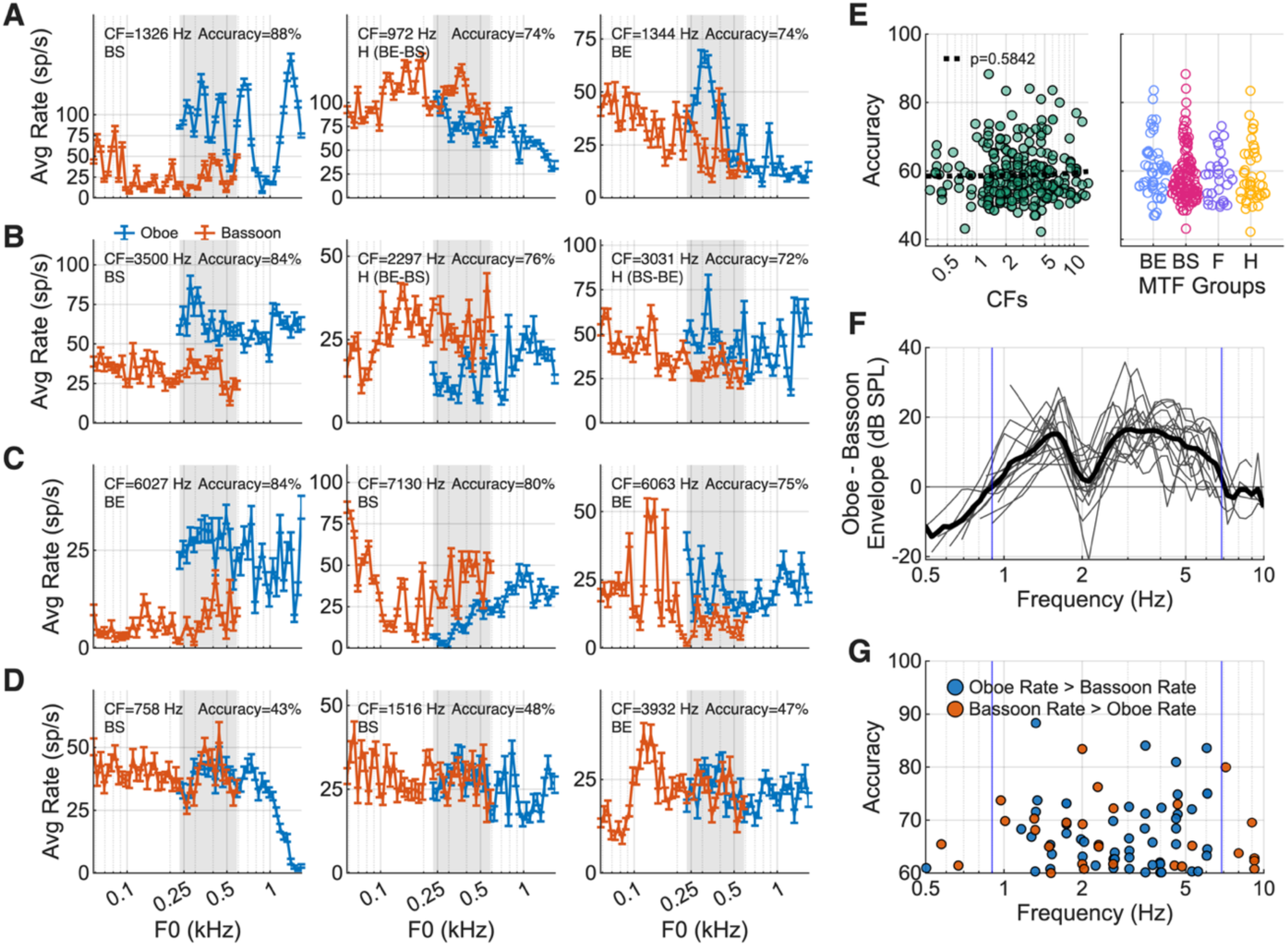
Example single-neuron average rate responses to oboe and bassoon and analysis of rate predictions of instrument identity. (A) Low-CF, (B) medium-CF, and (C) high-CF neuron examples of responses to oboe and bassoon. Grey bar indicates the range of overlapping F0s between oboe and bassoon. (D) Examples with poor instrument identification. (E) Accuracy of instrument identity classification for each neuron as a function of CF or classified by MTF type. (F) Differences between oboe and bassoon spectral envelopes (oboe – bassoon) (blue vertical lines indicate zero crossings of mean differences). (G) Prediction accuracy as a function of CF split into two groups, oboe rate greater than bassoon (blue), and bassoon rate greater than oboe (orange).

The difference in the spectral envelopes for bassoon and oboe over the 16 stimuli with overlapping F0 values was calculated to determine whether high identification accuracy in neurons could be explained by energy differences between the stimuli (Fig. 3f). Oboe stimuli had higher spectral magnitudes than bassoon from 900 to 6875 Hz with a dip near 2 kHz (Fig. 3f). If energy were the sole driver of the neural response rates, then average rates in response to oboe would have been greater than for bassoon from 900 to 6875 Hz. However, the results were mixed: rates in response to bassoon were often greater than for oboe even in the frequency region where oboe spectral envelopes had larger magnitudes, and accuracy did not depend on spectral magnitude, indicating that energy was not the sole driver of these responses (Fig. 3g). This result also raised the question whether temporal features, rather than energy differences, contributed to accurate instrument identification.

Next, we tested whether an SVM model using timing information (PSTHs) could classify instrument identity. The PSTH bin size that resulted in the highest mean accuracy was 20 ms, though accuracy did not differ across bin sizes from 0.75–150 ms (ANOVA, Fig. 4a). The 20-ms bin width indicates that phase-locking information about F0 did not dominate the identification accuracy, and instead the differences in onset and sustained activity were used to classify the two instruments (Fig. 4b). The mean neural accuracy for this task was 62%, with a range of 43 to 91%. For 70% of neurons, classification based on timing information was better than for average rates (Fig. 4c). Accuracy was correlated with CF (p=0.0463, linear regression) but did not differ across MTF groups (p=0.1431, Kruskal-Wallis).

**Figure 4.**
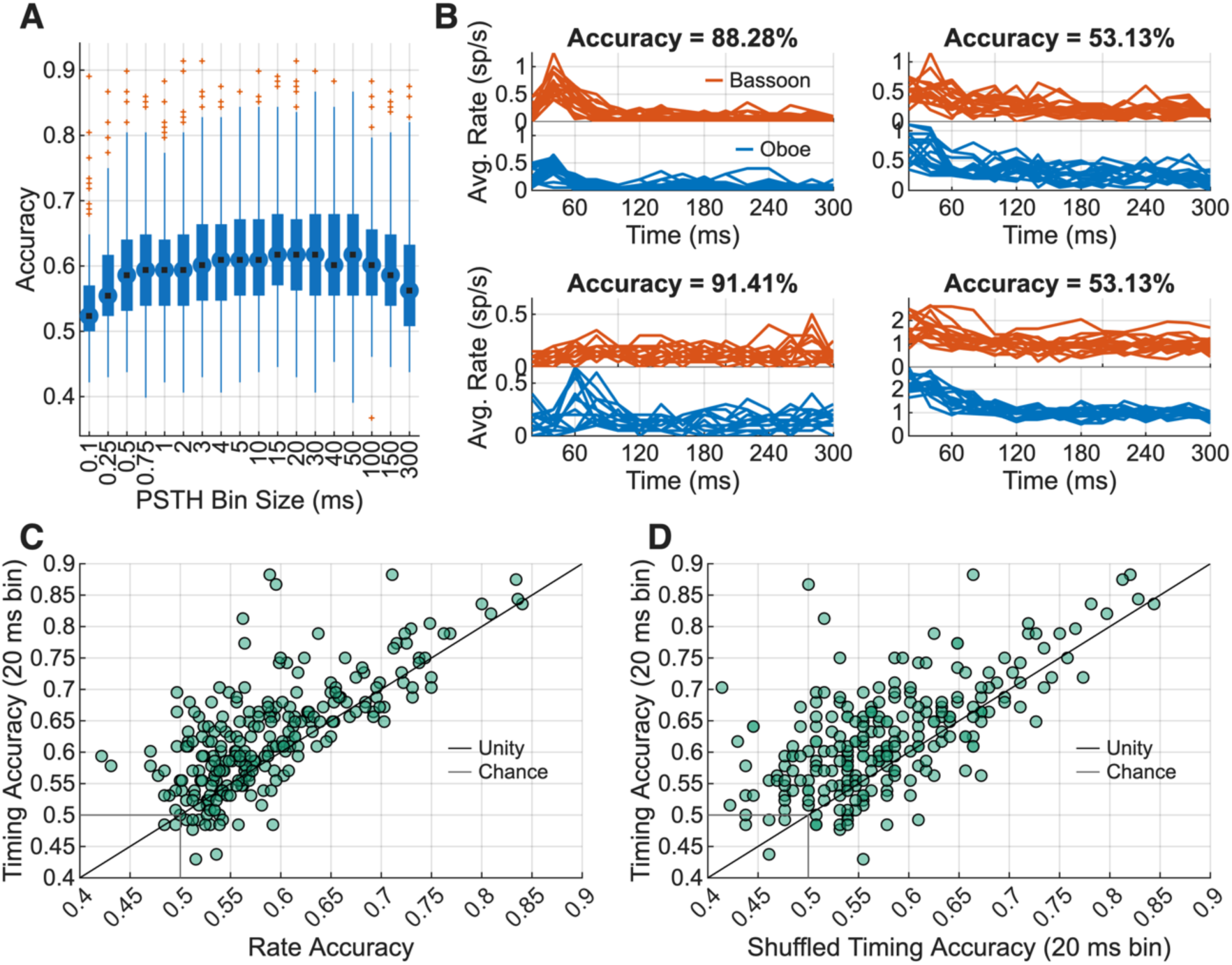
Single-unit instrument identification based on timing information. (A) Accuracy for all neurons as a function of PSTH bin size (outliers, red; median, black). (B) Example PSTHs with 20-ms bins for four example neurons, oboe (blue) and bassoon (orange) responses for each repetition. Top left: CF = 5278 Hz, BE. Bottom left: CF = 10556 Hz, Hybrid. Top right: CF = 2297 Hz, BS. Bottom right: CF = 2633 Hz, BS. (C) Timing accuracy plotted for each neuron as a function of rate accuracy. (D) Comparison of timing results without (vertical) and with (horizontal) shuffled PSTH timing, with a 20-ms bin widths, n=237 neurons.

Because timing-based identification outperformed rate-based identification, we investigated how much of this advantage depended on the specific temporal structure of the response by shuffling the PSTH bins within each trial. Shuffling the PSTHs reduced the accuracy (p<0.001, Mann-Whitney U-test), indicating that temporal structure was necessary for accurate instrument identification (Fig. 4d). However, this temporal structure includes both onset dynamics and F0-related periodicity and it is possible that the latter added noise rather than useful information when discriminating instruments across a range of F0s.

#### Instrument identification using rate and timing information across a population

An SVM classifier was trained on average-rate responses for a population of neurons to determine how a population of IC neurons encoded instrument identity. The population of CFs spanned 0.3–13.9 kHz, covering the range of the stimulus spectrum (Fig. 5a). When comparing driven rates in response to an example F0, overall population responses were similar but had small differences (Fig. 5a). Identification accuracy using the average rates of all neurons was 100%, compared to a distribution of chance-level accuracies obtained by fully randomizing rate values across neurons and trials while holding instrument labels fixed (100 permutations, Fig. 5b). To investigate the contributions of each neuron in the model, feature weights, called beta weights, were plotted as a function of CF (regression, p=0.4855), or classified by MTF group (p=0.0744, Kruskal-Wallis, Fig. 5c, d). A correlation between beta weights and single-unit rate accuracy was found (p<0.001, regression, Fig. 5e). A permutation test, in which a single neuron was taken out of the model, was used to test the impact of a single neuron on performance. The test revealed that removing any single neuron did not significantly reduce model performance (not shown), indicating that instrument-identification information was encoded redundantly in many neurons. To further investigate contributions of CF and MTF groups, models were trained using 1 to 100 neurons, sorted by beta weights. The model using the highest weighted neurons reached 99% accuracy with 5 neurons and 100% accuracy with 20 neurons (Fig. 5f, g, green). When neurons were split by CF group and the model was re-trained, the low-CF and medium-CF groups reached 99% accuracy when 80 or 75 neurons were included, respectively (Fig. 5g). Similarly, when only BS neurons were included, a model with 50 neurons identified instrument identity with 99% accuracy (Fig. 5g). Overall, instrument identity was redundantly encoded in many neurons, but a diverse set of neurons was needed to identify instruments using a small number of neurons.

**Figure 5.**
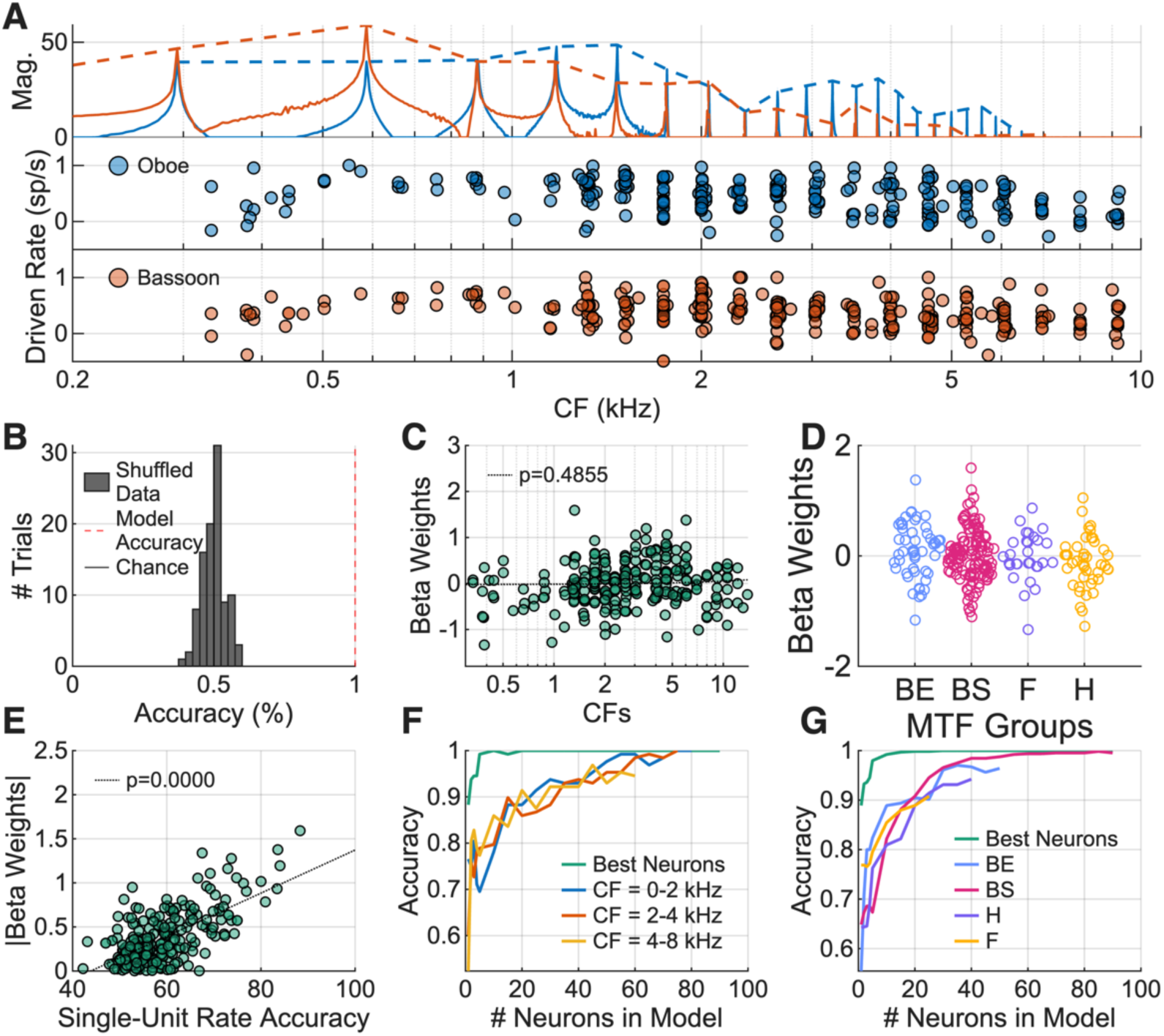
Instrument identification task using a population of neural rates. (A) Example population rate responses to a single F0 in either bassoon or oboe. (B) Decoding model performance compared to a null distribution of shuffled performance, generated by randomizing rate values across neurons and trials while holding instrument labels fixed (100 permutations) (C) Neuron weights plotted as a function of CF. (D) Neuron weights plotted by MTF group. (E) Single-neuron weights in the model as a function of the single-unit rate accuracy. (F) Model performance when using the highest weighted neurons for all neurons, low-, medium-, or high-CF neurons. (G) Same as F, but with MTF types.

Having established that rate information was sufficient to identify instrument identity at a population level, we next tested whether timing information showed a similar pattern. These results were similar to the rate results: 10 neurons diverse in CF and MTF type yielded 99% accuracy (Fig. S1). Overall, results are consistent with previous spectral-discrimination tasks using synthetic stimuli: both rate and timing responses encoded spectral information about the instrument identity.

### F0 Identification

#### F0 identification for oboe and bassoon using rate or timing information in single neurons

Next, the stimulus F0 of bassoon, oboe, or both, was identified based on single-neuron responses using an SVM to determine how F0 information was represented in these responses. The rate of a single neuron was a poor predictor of F0 identification for both bassoon and oboe, with mean identification accuracies of 6%, and highest accuracy of 17% and 16% (Fig. 6a, b). F0-identification models using timing information were more accurate: mean accuracy for identifying bassoon F0s was 14% with a maximum accuracy of 61%, and mean accuracy for identifying oboe was 6% and a maximum accuracy of 17% (Fig. 6a, b). F0-identification results based on responses of single neurons to bassoon and oboe were correlated: neurons with responses that allowed accurate identification of F0 in bassoon stimuli also contained information for accurate F0 identification in oboe F0 (p<0.001, regression, Fig. 6g). We further investigated the timing information present in the F0-decoding task for responses to bassoon. There was no correlation between accuracy and CF (p=0.3597, regression). BE neurons were significantly more accurate than other MTFs (p=0.0019, Kruskal-Wallis), and there was a correlation between RVF PCA 2 score and accuracy (p= 0.0094, regression, Fig. 6c, d, e). The highest decoding accuracy occurred at low F0s; decoding accuracy from the top 15 neurons was over 50% for F0s 58 to 196 Hz (Fig. 6f). The reliability of spike timing was highly correlated with the F0-identification accuracy for responses to bassoon, but vector strength was not (Fig. 6f, h, i). These results suggest that the temporal responses of neurons contained information about lower F0s. Reliability was also a better predictor of F0-identification accuracy than vector strength to F0, suggesting that though vector strength may be low, precise timing in IC neurons can be decoded.

**Figure 6.**
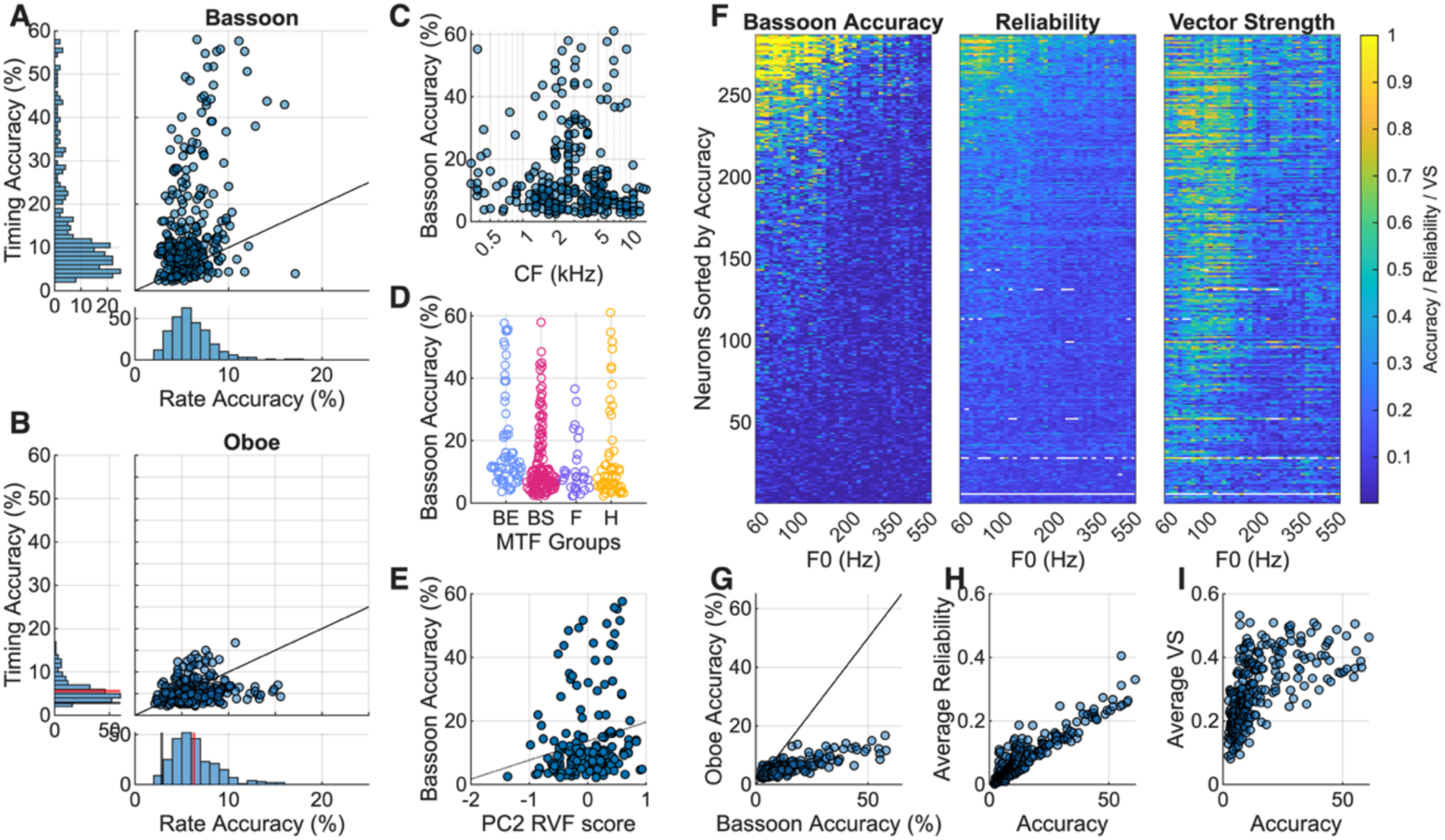
Decoding F0 based on single-neuron rate and timing information.(A) F0-identification accuracy based on timing for each neuron as a function of accuracy based on rate, for bassoon stimuli and (B) oboe stimuli. (C) F0-identification accuracy based on timing in response to bassoon as a function of CF, (D) MTF, or (E) PCA 2 from the RVF of the responses. (F) Accurate predictions for each bassoon F0 for all neurons, sorted by highest accuracy (left). Reliability (middle) and vector strength (right) calculations for each F0 and each neuron also sorted by highest accuracy. (G) Oboe timing F0-identification accuracy vs. bassoon accuracy. (H) Average reliability as a function of F0-identification accuracy for each neuron. (I) Average vector strength as a function of F0-identification accuracy for each neuron.

#### Neurons with accurate F0 identification contain reliable timing information

Neurons with reliable timing information resulted in more accurate F0 classifications, but reliability was only weakly correlated with vector strength. One example neuron with precise timing had a CF = 1529 Hz, BS MTF, and was sensitive to positive frequency sweeps (Fig. 7a, b, c). This neuron’s response was significantly phase locked to most bassoon stimuli (Fig. 7d). Period histograms revealed that, for most bassoon stimuli, the neuron fired at the same phase of the stimulus, but there were often two spikes per period (Fig. 7e).

**Figure 7.**
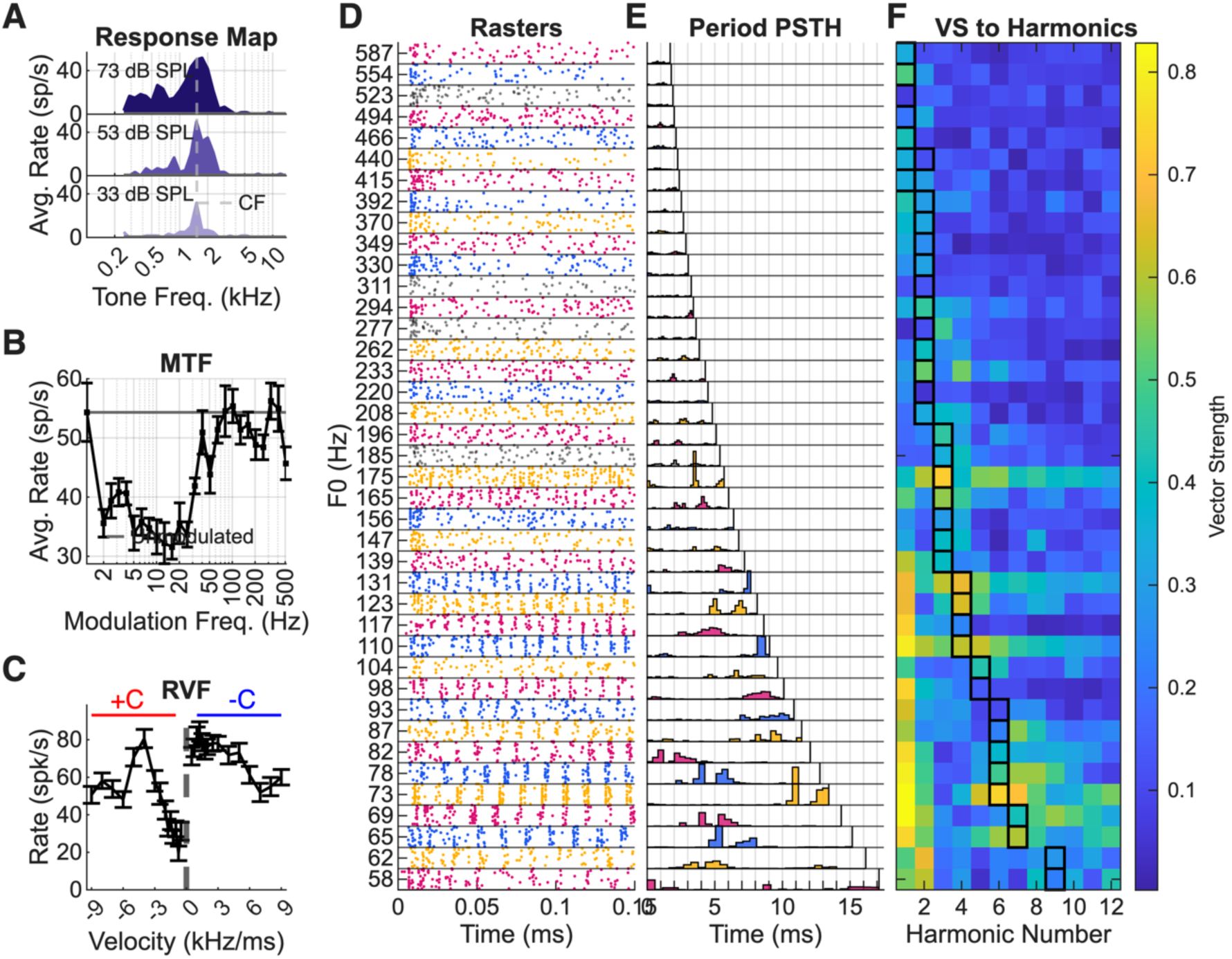
Neuron with the highest F0-identification accuracy. Response map, (B) MTF, and (C) RVF of the neuron. (D) PSTH and (E) period histogram of the neural responses to different bassoon F0s reveals complex responses. (F) Vector strength plotted for each harmonic, with the peak harmonic of each stimulus outlined in black.

Calculating vector strength to each harmonic revealed that this neuron had strong phase-locking responses to the fundamental but also exhibited phase locking near or at the spectral peak harmonic, which was approximately 500 Hz for all stimuli (Fig. 7f). This pattern of phase-locking near the peak harmonic was found in multiple neurons with high accuracy; neurons that had multiple spikes per period often phase-locked to the peak harmonic (Fig. 7, Fig. S2).

#### F0 identification for oboe and bassoon using rate or timing information in a population

F0 was identified using SVM classification models, which selected among the closed set of stimulus F0s based on rate and timing information from populations of neurons. Linear regression models were also used to predict F0 as a continuous value from the same population responses. First, an SVM classifier was used to identify bassoon or oboe F0 based on the rate responses of the entire population of recordings and reached 100% and 99% accuracy respectively (Fig. 8a). We also tested whether a simpler, linear regression model could predict F0. Linear regression models performed well, with R^2^ between the F0 predicted by the model and the actual F0 of 0.97 and 0.96 for both tasks (Fig. 8b). F0 was predicted within 5 semitones for bassoon and oboe (Fig. 8b). Beta weights varied with CF for the oboe (p<0.001). We split up neurons into CF groups and MTF groups and tested model accuracy using those groups against using all neurons, using the highest beta-weight neurons in the model first (Fig. 8c, d). Similar to results in the instrument-identification task, responses of a small number of diverse neurons yielded accurate F0 identification (Fig. 8c, d). Models based on neurons from a single CF or MTF group had worse performance. Interestingly, a linear readout of F0 using rate responses may exist, but it was necessary that the neurons included in the regression model were diverse in CF and MTF group.

**Figure 8.**
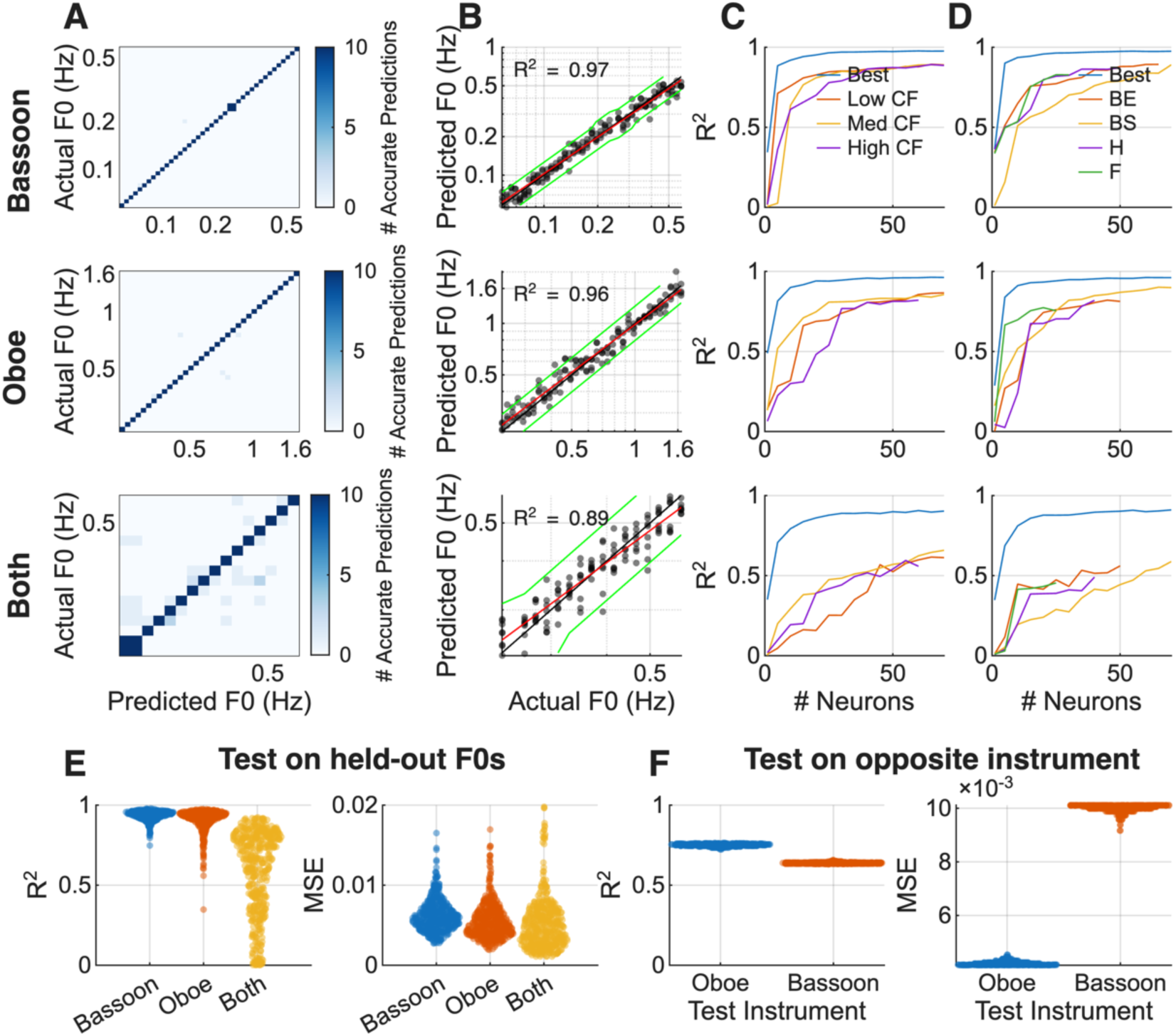
F0 identification based on rate responses for a population of neurons. (A) Confusion matrices for classification model. (B) Linear regression model predictions, fit line (red) and lines indicating 5 semitones above/below F0 (green). (C) Accuracy of a linear regression model using a subset of highest beta-weight neurons split into CF groups or (D) MTF groups. (E) Results of training on 80% of F0s and testing on 20% of F0s for bassoon, oboe, and both conditions, R^2^ and mean squared error results, 500 trials. (F) Results when linear regression model was trained on bassoon and tested on oboe (blue) and when model was trained on oboe and tested on bassoon (orange), 500 trials.

Though this task was framed as an F0-identification task, spectral centroid also varied as a function of F0 (Fig. 1). Thus, these models could be using information present in the responses related to F0 and/or to spectral centroid. To specifically investigate F0 information in these neurons, an SVM classifier was used to identify F0 and a linear regression model was used to predict F0, regardless of instrument identity. The stimuli included F0s from 233–587 Hz, which was the F0 range over which stimuli for both instruments were available. SVM performance reached 95% accuracy, and the R^2^ for the linear regression model was 0.89 (Fig. 8a, b). Beta weights were correlated with CF (p=0.0014). Linear regression models were run with a subset of neurons sorted by beta weights and split into CF groups or MTF types (Fig. 8c, d). Using all MTF and CF groups, models reached a maximum R^2^ of 0.90 using 50 neurons (Fig. 8c, d). However, performance decreased when the model included only neurons from single MTF or CF groups. The overall drop in performance for this task compared to the bassoon or oboe tasks indicated that the neural responses to the instrument spectrum had an impact on F0 identification.

Next, we tested how well linear-regression models generalized for two tasks: 1) training on a subset of F0s and using the model to predict new F0s (Fig. 8e), and 2) training F0 prediction using responses to one instrument and testing with responses to the other instrument (Fig. 8f). Testing on held-out F0s resulted in R^2^ values near those for the original F0-identification tasks (Fig. 8b), for which bassoon F0 prediction had a mean R^2^ of 0.94, oboe F0 prediction had a mean R^2^ of 0.91, and predictions of F0s for both instruments had a mean R^2^ of 0.58 (Fig. 8e). For the second task, the mean R^2^ for models trained on bassoon data and tested on oboe F0s was 0.75, and interestingly, the models trained on oboe F0s and tested on bassoon F0s had an R^2^ of 0.64 (Fig. 8f). These results suggest that the models generalize to different F0s, but there are limitations in generalizing across different instruments.

Next, SVM models were used to identify F0 based on timing information from a population of neurons. Again, F0 was identified for bassoon and oboe stimuli separately, and then F0s were identified for oboe and bassoon for overlapping F0s. Models were run using 1 to 200 neurons in the population, sorted by individual-neuron timing accuracy. Bassoon F0 identification rose to 99% accuracy with 30 neurons (Fig. 9a, top). Oboe F0 identification results had similar trends, though 100 neurons were required to reach 88% accuracy (Fig. 9a, middle). For both tasks, higher F0s were more accurately classified as more neurons were added to the model (Fig. 9b, top, middle). These results indicated that F0 and spectral changes with F0 are also encoded in neural timing information in lower-F0 regions.

**Figure 9.**
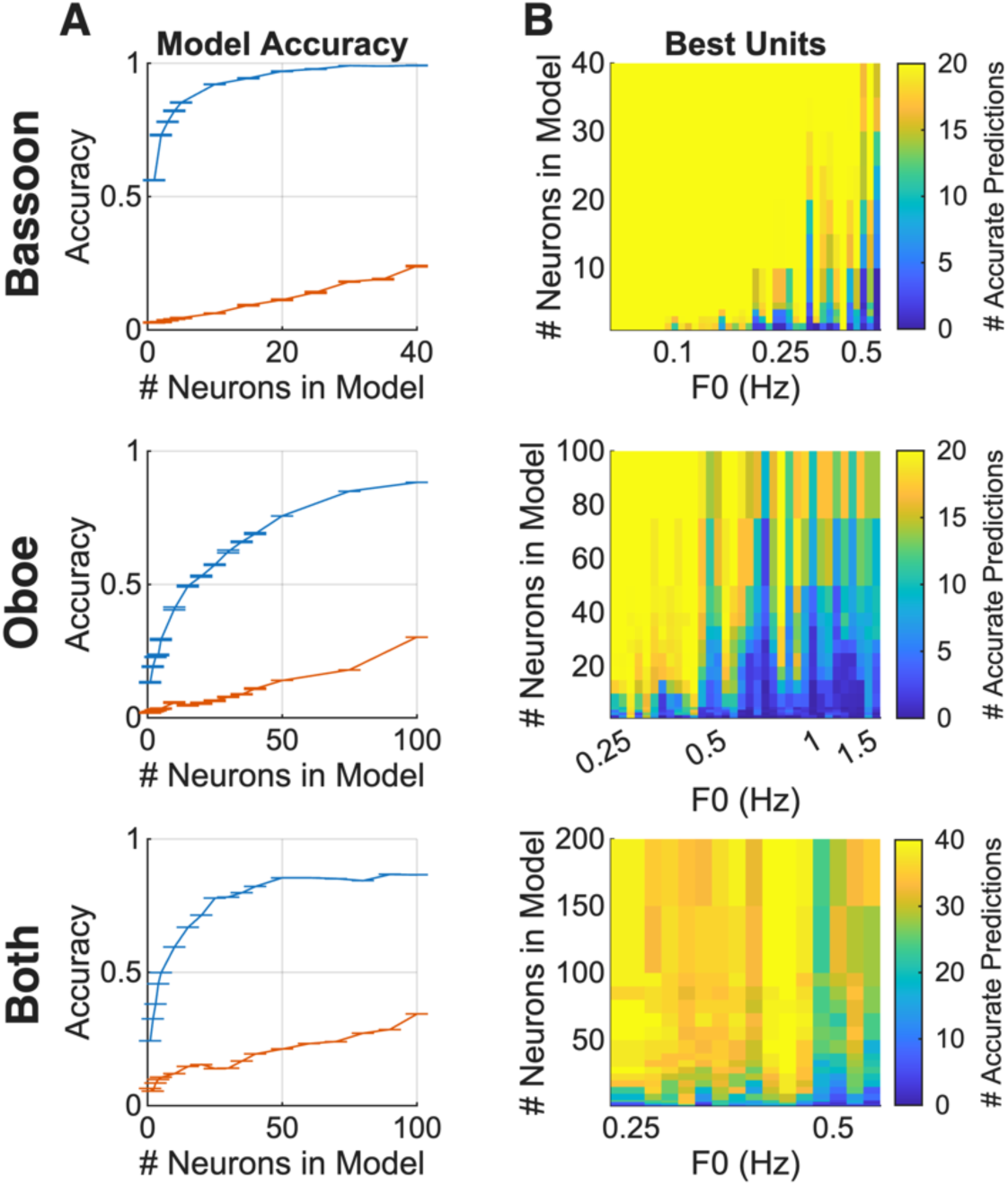
F0-identification accuracy using multiple neuron PSTHs for bassoon, oboe, and both bassoon and oboe. (A) F0-identification accuracy as a function of number of neurons in the model, with neurons sorted by best (blue) or worst (red) based on individual neuron PSTH F0-identification accuracy. (B) Number of accurate identifications for each F0 as the number of neurons in the model increases.

The F0-identification model based on timing in responses to bassoon and oboe for the range of overlapping F0s was created to test for F0 identification while avoiding differences due to spectral centroid. The SVM model using this information reached 88% accuracy based on 200 neurons (Fig. 9a, bottom). F0 identification was less accurate over all F0s when spectrum information was mixed (Fig. 9b, bottom). Overall, spectral changes due to instrument identity across F0 values aided in identification accuracy in this neural population for analyses based on both timing and rate information.

#### Instrument/F0 Identification

The result that spectral information improved F0 identification in our models led us to investigate whether information about both F0 and instrument identity was encoded by the same neurons. The bassoon F0-identification and instrument-identification accuracy based on single-neuron timing information had the overall highest accuracies and were positively correlated (bassoon rate p<0.001, bassoon timing p<0.001, oboe rate p<0.001, oboe timing p<0.001, Fig. 10a, bottom left). In fact, bassoon or oboe F0-identification were correlated with instrument-identification accuracies for both rate and timing of single-neuron responses, though accuracies were overall poor (Fig. 10a).

**Figure 10.**
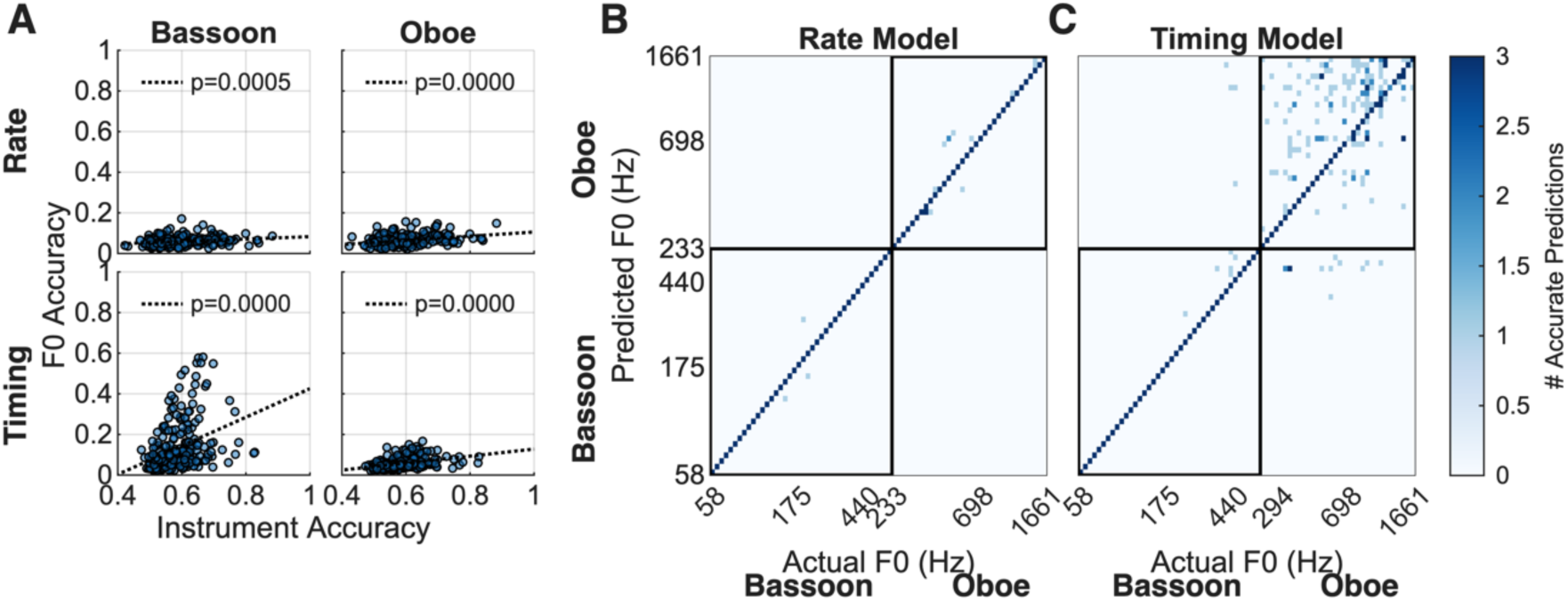
Investigating pitch/timbre interference. Scatter plots of accuracy of F0 identification for specific instruments based on response rate or timing for bassoon and oboe F0s. (B) Confusion matrix for the population model based on response rate for specific-instrument F0 identification. (C) Confusion matrix using timing information for 40 neurons.

Next, we used an SVM to decode both instrument identity and F0 for all 75 possible stimuli (Fig. 10b, c, d). This ‘token’ decoding task asked whether rate or timing were sufficient to decode specific instrument F0s in a population of responses. SVM identification accuracy based on response rate was 98.8% using all 247 neurons, whereas specific-instrument F0 identification based on timing was 85.9% using 40 neurons, at which point accuracy plateaued (Fig. 10b, c). The model based on timing information in the population responses again struggled with high F0s, confusing instrument identity and F0s at higher F0s (Fig. 10d). Within the population of neurons recorded here, enough information existed to decode both instrument identity and F0. Decoding based on response rate, specifically, did not confuse instrument identity, whereas decoding based on response timing was less accurate and did confuse instrument identity. Over this population of IC neurons, rate responses were more robust than timing for instrument-specific F0 identification.

#### Why Current IC Models Fail

For each BE and BS neuron in the neural population, we simulated responses using an SFIE model with the CF, MTF shape, and best/worst modulation frequency matched to the given neuron. Response map and MTF were used to determine the SFIE model parameters, and bassoon and oboe stimuli were used as inputs. Model responses were also calculated using the BMSI model and an energy model. We analyzed average rate responses to bassoon and oboe stimuli and compared the model rates with the neural rate responses. We found that the variance of the data explained by the model was at chance for most of the data (Fig. 11a). IC models did not accurately predict rate changes as F0 increased in bassoon or oboe.

**Figure 11.**
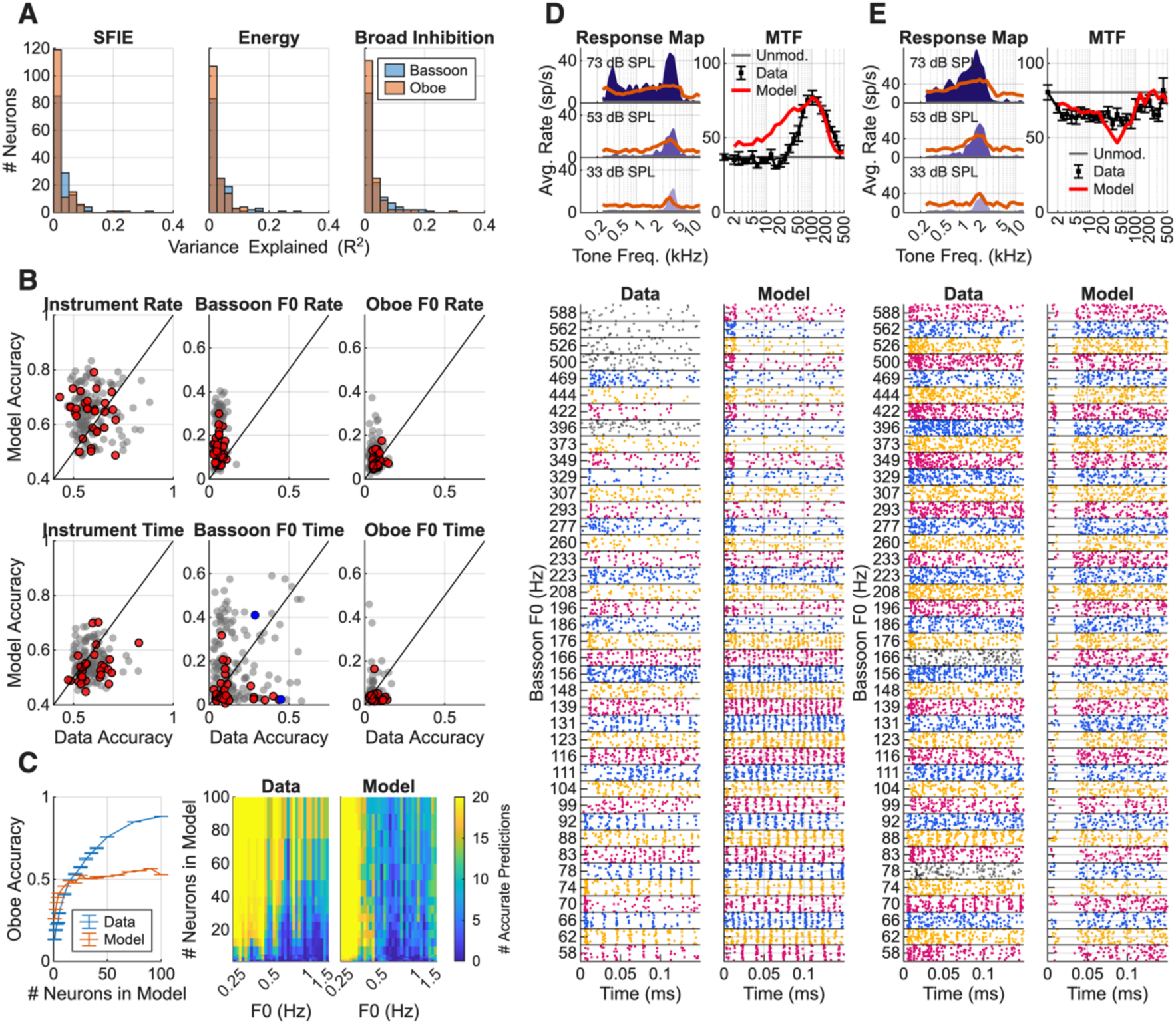
Model predictions of instrument and F0 identity and comparisons to neural data. (A) Variance of the neural rate responses to bassoon and oboe explained by the model responses, SFIE, energy, and BMSI for each neuron. (B) SFIE-model instrument-and F0-identification accuracy, plotted as a function of neural accuracy for each single neuron. Unity line (black). Grey dots indicate models with responses that explained less of the variance in the neural response maps and MTFs (R^2^ < 0.5), red dots indicate models that explained more of the variance in the data (R^2^ > 0.5). Blue dots indicate examples in (D) and (E). (C) Population decoding of oboe F0 for data and SFIE model, including overall accuracy as more neurons are added to the decoding model (left), and accuracies for each F0 prediction (middle data, right SFIE model). (D) Response map, MTF, and bassoon response dot rasters for data and model BE neuron and (E) BS neuron.

Next, we used the decoding models to investigate differences between SFIE model and neural responses. The spike times and average rates from the SFIE model were used in the same decoding tasks as the data. Decoding F0 or instrument identity in single model neurons often had similar distributions of accuracies as the data (Fig. 11b). However, there was no correlation between data accuracy and SFIE accuracy for single neurons, except in the oboe and bassoon F0-identification task based on response rates (p=0.0150, p=0.0126, Fig. 11b). However, we questioned whether the response maps and MTFs of the SFIE model accurately reflected neural response maps and MTFs. The correlation between the neural and SFIE model response maps and the correlation between neural and SFIE model MTFs were calculated. The correlations were squared and then averaged to estimate the neural variance explained by the SFIE model. Model and data accuracies were not significantly correlated.

To further investigate differences between the SFIE-model responses and the neural data, we compared temporal responses to bassoon and oboe (Fig. 11). For both example neurons, the R^2^ between neural and model MTFs and response-maps were above 0.5. Despite MTF and response-map similarity, temporal patterns for bassoon stimuli differed from those in neural data for both examples (Fig. 11d, e). The first example had an SFIE-model response that could be decoded with similar accuracy as the neural response, but there are clear differences in onsets and temporal patterns (Fig. 11d). The second example features an SFIE-model response with little phase locking to stimulus features and poor decoding accuracy (Fig. 11e).

Population instrument and F0-identification accuracies were similar between the SFIE model and neural data (not shown). Instrument-identification based on the population SFIE model also relied on a diverse set of CFs and MTF types to encode the instruments, but in general contained redundant information. The population SFIE-model F0-identification accuracy based on model rate responses was also similar to that of the neural decoders, with bassoon identification 100% accurate, oboe identification 95% accurate, and both oboe and bassoon F0 identification 94% accurate. Next, F0 identification using SFIE timing information was similar for the bassoon F0 task, with SFIE accuracy of 96% with 30 neurons. Interestingly, one difference between neural decoding and SFIE decoding was found in decoding oboe F0 identity. In the SFIE model, decoding accuracy increased to 50% with 25 neurons in the model but additional model neurons did not improve accuracy, unlike in the data (Fig. 11c, left). Specifically, prediction accuracy sharply fell off at F0s greater than 370 Hz (Fig. 11c).

## Discussion

We recorded extracellular responses in the IC in awake Dutch-belted rabbits to bassoon and oboe sounds over a range of F0. Decoding models were used to characterize how instrument and F0 information were represented in single neurons and population responses. Information contained in rate responses of a population was sufficient to identify F0 and/or instrument identity and models had the best performance when neurons were diverse in CF and MTF type. Additionally, decoding models could use timing information to identify F0 and instruments. Models used information from single-neuron timing responses to identify lower F0s. Population timing information also identified F0s, though accuracy decreased as F0 increased.

### Rate responses were sufficient for timbre encoding in quiet

Single neurons contained rate and timing information sufficient to identify instruments with up to 88% accuracy. This result differed from a study using recorded vowels, in which rate-based vowel identification was poor, but timing-based vowel identification was accurate [6]. This difference could be due to the variation of F0 information in the instrument-identification task and the higher F0s used, whereas the previous study identified sets of vowels at a single F0, ranging from 95–202 Hz [6]. It is likely that more precise timing in neurons at lower F0s contributed to higher decoding accuracy based on response timing in the vowel study.

Rate or timing information from a population of IC neurons could also be used to identify instrument identity. Previously, behavioral spectral-peak discrimination thresholds in quiet could be explained by rate responses in budgerigar IC, and human spectral-peak discrimination thresholds were similar to neural discrimination thresholds in rabbit IC rate responses [1, 5]. Interestingly, temporal information was necessary to discriminate spectral peaks in noise in budgerigar IC, providing a possible role for the redundant encoding of spectral information using timing [5]. We also found that neurons that were diverse in CF and MTF types yielded the best model performance with the least number of neurons.

A critical area of timbre investigation is identifying which stimulus attributes are essential for instrument identification. The presence of frequency sweeps in natural stimuli may affect encoding: a vowel study in the IC found that decoding accuracy based on response timing was significantly correlated with the second principal component of the RVF [6]. Psychophysical studies have examined correlations between perceptual attributes of timbre to specific sound characteristics (Review: [21]), and it is unknown how most attributes are encoded in the IC. A recent study found that spectral peaks, which are related to the spectral centroid and the percept of brightness, are encoded in the IC [1]. However, other attributes most likely also contribute to timbre encoding, for example, attack time could largely impact temporal responses, enhancing temporal information about instrument identity. Though the significance of these attributes could not be tested in the present study, which focused on steady-state sounds, future studies could systematically vary these attributes to further investigate this question.

### Both rate and timing in responses contained information about F0

In responses of single neurons, decoding models were more accurate when trained on timing information compared to rate information. One study of IC responses to HTCs found that rate information could reliably encode F0s > 800 Hz [3], but we found no evidence of an increase in accuracy for high F0s in models trained on rate responses (Fig. S3). Phase locking was present up to 900 Hz in HTCs [2]. Neurons in our study significantly phase-locked to F0 up to approximately 600 Hz, but generally, decoding models could only use temporal information up to 196 Hz. Differences between the HTCs and instrument results could be due to differences in the amplitude of harmonics, but more work needs to be done to examine differences between encoding F0 in flat and non-flat spectrums.

Decoding models trained on population information were much more accurate than single neurons, as expected. F0 identification using models trained on rate information from a population was accurate with either an SVM model or linear regression. The linear regression model relied on a small subset of neurons with diverse CFs and MTF types, and rate responses in each of these neurons had a relatively linear relationship with F0. The linear model was chosen over the SVM to test the simpler hypothesis that F0 can be decoded linearly from rate. Model results were constant as a function of F0, supporting the hypothesis that rate can encode higher F0s, where responses to individual components may be resolved [2], but interestingly lower F0s were encoded in rate responses as well. Note, these studies were all performed in rabbit; changes in tuning for different animal species would likely impact these results.

F0 identification trained on timing information for a population of neurons also had high accuracy. However, at higher F0s, temporal decoding models struggled, even as more neurons were added to the model. This decline in accuracy at higher F0s likely reflects a fundamental limit on the precision of phase-locking in the IC: as F0 increases, the auditory system’s ability to phase-lock to individual stimulus cycles degrades, reducing the temporal information available to the population regardless of how many neurons are included in the model. This finding is consistent with the phase-locking limits reported in previous work using synthetic harmonic complex tones (up to 900 Hz).

### Pitch-timbre interference may be present in the IC

This study investigated whether pitch and timbre interference was present in the IC. Interference was investigated by using decoding models that 1) predicted F0 across a single instrument or 2) predicted F0 across both instruments for the F0s where oboe and bassoon overlapped. Decoding F0 in both instruments decreased model performance when models were trained on both rate and timing population responses. This decrease in performance indicates that the decoding model was taking advantage of spectral changes due to F0 in the single-instrument F0 identification task. Additionally, single neuron accuracy of F0 identification was highly correlated with instrument identification accuracy, indicating that neurons often encode both pitch and timbre. These results support the hypothesis that pitch and timbre information are jointly encoded in the IC, and that variation in timbre disrupts pitch discrimination in IC neurons [10].

### Benefits and limitations of computational modeling approaches

Decoding models can be used to reveal neural representations of information in a brain region, but results must be interpreted carefully (Review: [22]). First, though decoding models are useful for finding information in IC responses, it is unknown whether or how downstream auditory structures actually extract and use this information for perception or behavior; the presence of decodable information in IC responses does not establish that the brain performs an equivalent readout. Decoding models are not biologically plausible, in general. The SVM and linear regression models used here combine information across neurons using weighted sums optimized to maximize decoding accuracy, a computation with no known biological analogue. There is no evidence that the medial geniculate nucleus, or any other downstream structure, combines IC inputs in this specific way. Second, weights of decoders are difficult to interpret, due to the nonlinear combinations in the SVMs.

Another approach for estimating each neuron’s contribution to the decoding model is a leave-one-out permutation test: each neuron is individually removed from the population, the model is retrained on the remaining neurons, and the resulting change in model accuracy indicates that neuron’s contribution to overall performance. In our data, removing one neuron did not alter performance, generally, due to the redundant information present in the IC, suggesting that no individual neuron was critical to model performance. Instead, we trained models on subsets of neurons to determine what types of neurons carried important pitch or timbre information. Lastly, model overfitting to a given dataset can lead to high identification accuracies even when information is not present in the neural data. We used 5-fold cross validation to prevent overfitting, but also analyzed neural responses used in the models to validate the presence of information.

Other computational IC models have been used to investigate specific encoding mechanisms in the IC. SFIE, BMSI, and energy models were poor predictors of bassoon and oboe rate responses over many F0s. However, other work has found that the SFIE-model predictions, with sensitivity to amplitude modulation and neural fluctuations, significantly correlated with a majority of neural responses to different vowels [7]. Building upon that work, a model that was sensitive to amplitude modulation and frequency-sweep direction/velocity better predicted vowel identification using timing information [6]. Separately, the SFIE model with the addition of BMSI more accurately modeled IC responses to spectral peaks of a harmonic stimulus [1]. Though the SFIE model was a poor predictor of average rate in response to bassoon and oboe F0s, we used decoding models with SFIE model inputs to examine how the information in the model differs from neural responses. We found that the model’s temporal response patterns differed substantially from the neural data, and that these models also produced average-rate responses that differed from the neural data. More work needs to be done to investigate these differences in depth.

### Human vs. Animal Pitch/Timbre Perception

It is important to consider differences between human and animal perception when studying pitch and timbre. It has been hypothesized that humans generally use rate information, because humans are better at pitch discrimination when resolved harmonics are present [23] and cortical responses are more effectively driven by resolved harmonics [24]. Most animal models, however, with broader peripheral tuning, are better at pitch tasks when harmonics are unresolved [25]. Previous behavioral work has found that ferrets, gerbils, and chinchillas may be more reliant on temporal cues, whereas marmosets rely on resolved harmonics [25–28]. However, recent behavioral studies suggest that rabbits can discriminate F0s without the fundamental in the ranges 200 – 400 Hz, 400 – 800 Hz, and 800 – 1600Hz using resolved harmonics at high F0s and timing at lower F0s [4].

The analyses here focused on information present in IC neurons that may be useful in instrument and F0 identification, or both, but remains limited by the stimuli used. A more diverse set of stimuli, including responses from instruments with strong F0s (such as trombone), unusual spectral envelopes (such as clarinets), and time-varying sounds (such as piano) would be useful to tease apart differences in neuronal coding of these aspects of sound. Another stimulus to add to this experiment would be the addition of simplified, synthetized instrument sounds. For example, the impact of phase on encoding could be determined by playing a natural-instrument sound followed by a synthesized sound with harmonics in sine phase. To investigate how timbre interferes with pitch, a flat-spectrum stimulus could be created using the same harmonic phase relationships as the natural instrument sound, isolating the contribution of phase from that of spectral shape. Comparing neural responses to this phase-matched flat-spectrum stimulus with responses to the natural instrument sound would help determine whether phase information alone, independent of spectral envelope, contributes to pitch-timbre interference. This paradigm would help answer questions about encoding that this initial study leaves open.

## Acknowledgements

This study was funded by NIH-R01-DC010813 and NIH-F31-DC020630-03. Thanks to Douglas Schwarz for assistance with hardware and software. Thanks to Braden Maxwell for feedback on analysis. Thanks to Kris Abrams for helping with the rabbit surgery and care. Thanks to Dr. Daniel Guest for feedback on machine-learning models.

## Notes

Supported by: NIH R01-DC010813, NIH F31-DC020630

### Competing Interest Statement

The authors have declared no competing interest.

## Bibliography

1. Fritzinger JB, Carney LH (2026) Timbre Encoding in the Inferior Colliculus. J Neurosci 46:e1104252026. 10.1523/JNEUROSCI.1104-25.2026

2. Su Y, Delgutte B (2019) Pitch of harmonic complex tones: rate and temporal coding of envelope repetition rate in inferior colliculus of unanesthetized rabbits. Journal of Neurophysiology 122:2468–2485. 10.1152/jn.00512.2019

3. Su Y, Delgutte B (2020) Robust Rate-Place Coding of Resolved Components in Harmonic and Inharmonic Complex Tones in Auditory Midbrain. J Neurosci 40:2080–2093. 10.1523/JNEUROSCI.2337-19.2020

4. Wagner JD, Gelman A, Hancock KE, et al (2022) Rabbits use both spectral and temporal cues to discriminate the fundamental frequency of harmonic complexes with missing fundamentals. Journal of Neurophysiology 127:290–312. 10.1152/jn.00366.2021

5. Henry KS, Abrams KS, Forst J, et al (2017) Midbrain Synchrony to Envelope Structure Supports Behavioral Sensitivity to Single-Formant Vowel-Like Sounds in Noise. JARO 18:165–181. 10.1007/s10162-016-0594-4

6. Mitchell PW, Carney LH (2025) Chirp Sensitivity and Vowel Coding in the Inferior Colliculus

7. Carney LH, Li T, McDonough JM (2015) Speech Coding in the Brain: Representation of Vowel Formants by Midbrain Neurons Tuned to Sound Fluctuations1,2,3. 12

8. Allen EJ, Oxenham AJ (2014) Symmetric interactions and interference between pitch and timbre. The Journal of the Acoustical Society of America 135:1371–1379. 10.1121/1.4863269

9. McPherson MJ, McDermott JH (2023) Relative pitch representations and invariance to timbre. Cognition 232:105327. 10.1016/j.cognition.2022.105327

10. Maxwell BN, Fritzinger JB, Carney LH (2020) Neural Mechanisms for Timbre: Spectral-Centroid Discrimination based on a Model of Midbrain Neurons. p 4

11. Siedenburg K, Jacobsen S, Reuter C (2021) Spectral envelope position and shape in sustained musical instrument sounds. The Journal of the Acoustical Society of America 149:3715–3726. 10.1121/10.0005088

12. Bizley JK, Walker KMM, Silverman BW, et al (2009) Interdependent Encoding of Pitch, Timbre, and Spatial Location in Auditory Cortex. Journal of Neuroscience 29:2064–2075. 10.1523/JNEUROSCI.4755-08.2009

13. Quiroga RQ, Nadasdy Z, Ben-Shaul Y (2004) Unsupervised Spike Detection and Sorting with Wavelets and Superparamagnetic Clustering. Neural Comput 16:1661–1687. 10.1162/089976604774201631

14. Schwarz DM, Zilany MSA, Skevington M, et al (2012) Semi-supervised spike sorting using pattern matching and a scaled Mahalanobis distance metric. J Neurosci Methods 206:120–131. 10.1016/j.jneumeth.2012.02.013

15. Mitchell PW, Henry KS, Carney LH (2023) Sensitivity to direction and velocity of fast frequency chirps in the inferior colliculus of awake rabbit. Hearing Research 440:108915. 10.1016/j.heares.2023.108915

16. Gai Y, Carney LH (2006) Temporal Measures and Neural Strategies for Detection of Tones in Noise Based on Responses in Anteroventral Cochlear Nucleus. Journal of Neurophysiology 96:2451–2464. 10.1152/jn.00471.2006

17. Fritzinger JB, Carney LH (2025) Mechanisms of Tone-in-Noise Encoding in the Inferior Colliculus. Journal of Neuroscience 45:. 10.1101/2025.03.07.642101

18. Zilany MSA, Bruce IC, Carney LH (2014) Updated parameters and expanded simulation options for a model of the auditory periphery. J Acoust Soc Am 135:283–286. 10.1121/1.4837815

19. Nelson PC, Carney LH (2004) A phenomenological model of peripheral and central neural responses to amplitude-modulated tones. J Acoust Soc Am 116:2173–2186. 10.1121/1.1784442

20. Carney LH, McDonough JM (2019) Nonlinear auditory models yield new insights into representations of vowels. Atten Percept Psychophys 81:1034–1046. 10.3758/s13414-018-01644-w

21. McAdams S (2019) The Perceptual Representation of Timbre. In: Siedenburg K, Saitis C, McAdams S, et al (eds) Timbre: Acoustics, Perception, and Cognition. Springer International Publishing, Cham, pp 23–57

22. Kriegeskorte N, Douglas PK (2019) Interpreting encoding and decoding models. Current Opinion in Neurobiology 55:167–179. 10.1016/j.conb.2019.04.002

23. Shackleton TM, Carlyon RP (1994) The role of resolved and unresolved harmonics in pitch perception and frequency modulation discrimination. The Journal of the Acoustical Society of America 95:3529–3540. 10.1121/1.409970

24. Norman-Haignere S, Kanwisher N, McDermott JH (2013) Cortical Pitch Regions in Humans Respond Primarily to Resolved Harmonics and Are Located in Specific Tonotopic Regions of Anterior Auditory Cortex. J Neurosci 33:19451–19469. 10.1523/jneurosci.2880-13.2013

25. Walker KM, Gonzalez R, Kang JZ, et al (2019) Across-species differences in pitch perception are consistent with differences in cochlear filtering. eLife 8:e41626. 10.7554/eLife.41626

26. Shofner WP (2002) Perception of the periodicity strength of complex sounds by the chinchilla. Hearing Research 173:69–81. 10.1016/s0378-5955(02)00612-3

27. Klinge A, Klump GM (2009) Frequency difference limens of pure tones and harmonics within complex stimuli in Mongolian gerbils and humans. The Journal of the Acoustical Society of America 125:304–314. 10.1121/1.3021315

28. Song X, Osmanski MS, Guo Y, Wang X (2016) Complex pitch perception mechanisms are shared by humans and a New World monkey. Proc Natl Acad Sci USA 113:781–786. 10.1073/pnas.1516120113

