## Supplemental Figures for "Representations of Pitch and Timbre of Instrument Sounds in the Inferior Colliculus"

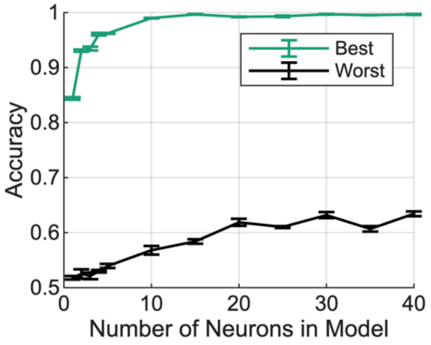


**Figure S1.** Decoding instrument identity using timing information in a population of neurons. Neurons were added to the model from best to worst (green) or worst to best (grey) sorted by single neuron accuracy in the decoding task using timing.

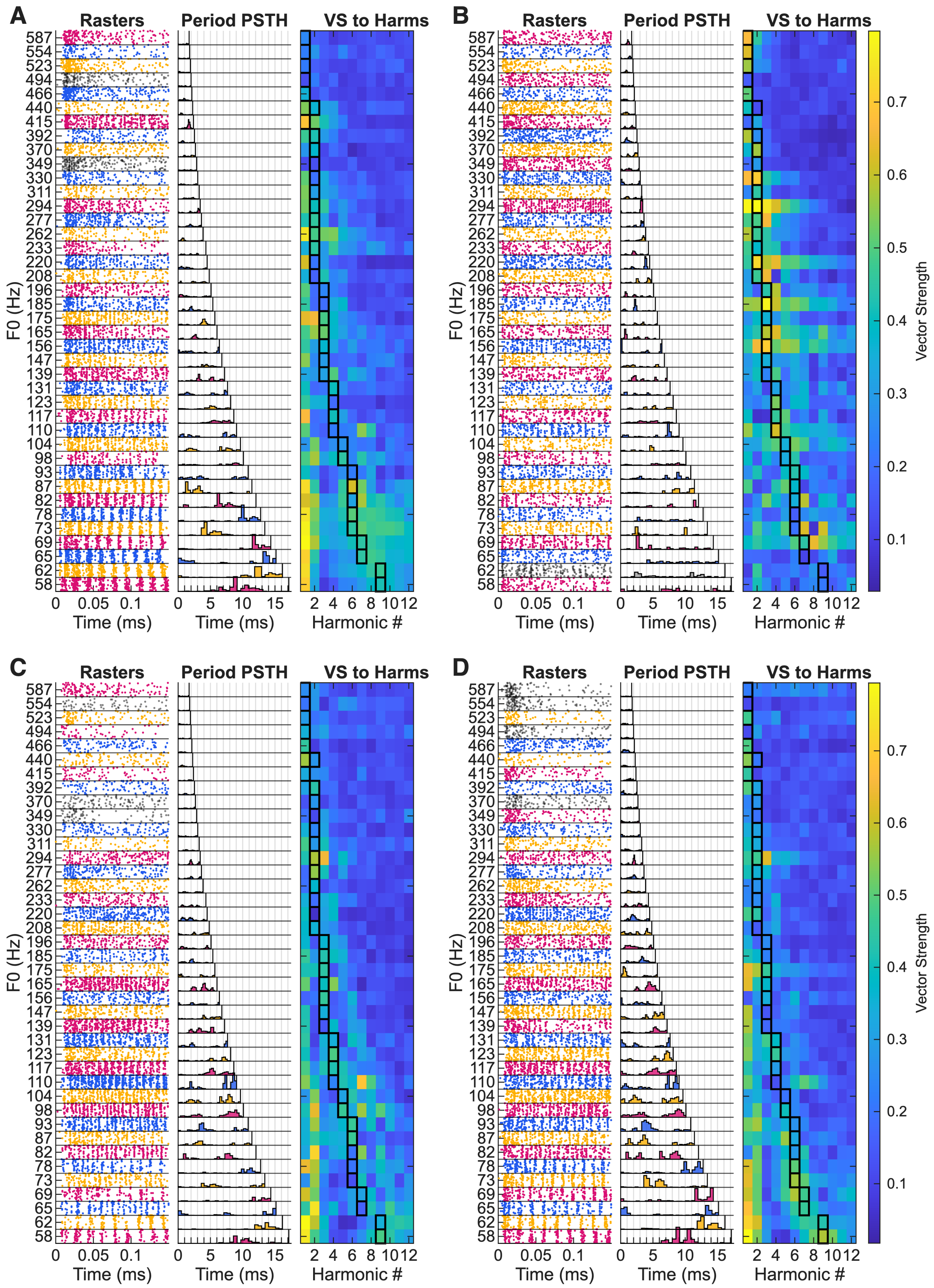
A
B
C
D
Rasters
Period PSTH
VS to Harms
F0 (Hz)
587
554
523
494
466
440
415
392
370
349
330
311
294
277
262
233
220
208
196
185
175
165
156
147
139
131
123
117
110
104
98
93
87
82
78
73
69
65
62
58
0
0.05
0.1
Time (ms)
0
5
10
15
Time (ms)
2
4
6
8
10
12
Harmonic #
Vector Strength
0.1
0.2
0.3
0.4
0.5
0.6
0.7

**Figure S2.** Four neuron temporal responses to bassoon stimuli. Dot rasters (left), period histogram (middle) and vector strength to the first 12 harmonics (right), grey rasters indicate non-significant phase locking to the F0. Black squares indicate peak harmonic of each stimulus. (A) CF = 6966 Hz, Hybrid. (B) CF = 2629 Hz, BS. (C) 6063 Hz, BE. (D) 3482 Hz, BE.

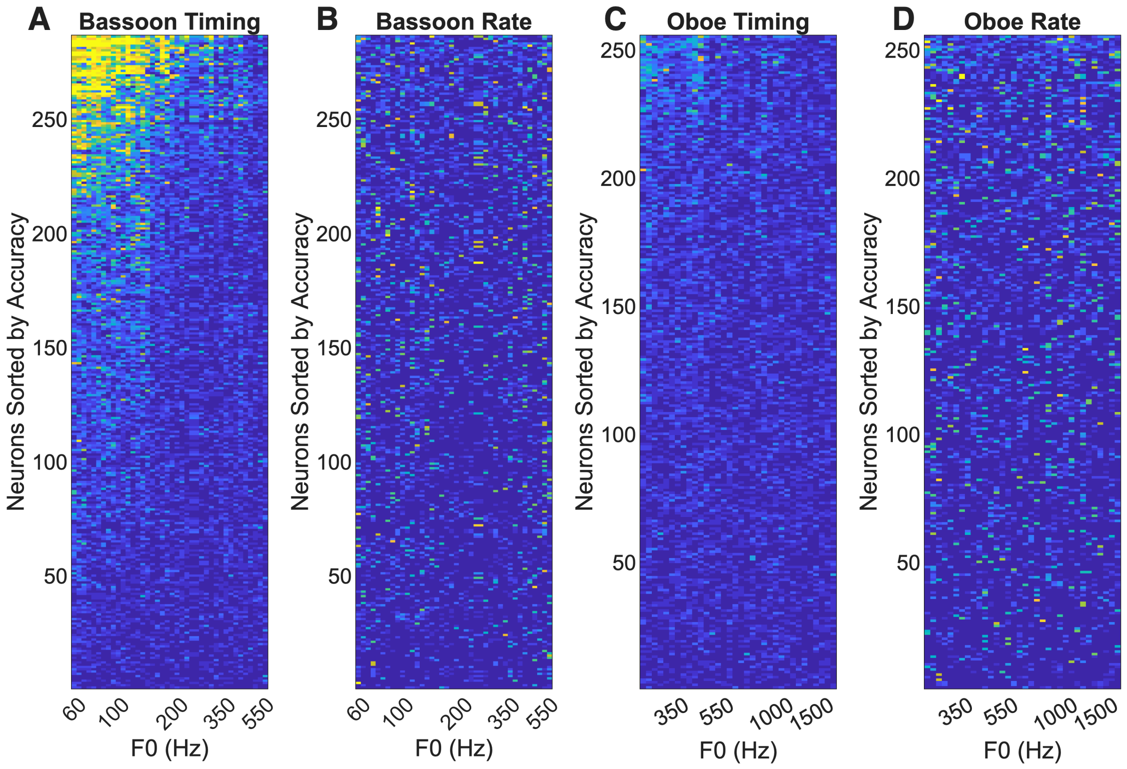


**Figure S3.** Decoding accuracy for each F0 for all neurons. Sorted so that the most accurate decoding of F0 is at the top. (A) Decoding results for bassoon F0 identification using timing. (B) Decoding results for bassoon F0 identification using rate. (C) Decoding results for oboe F0 identification using timing. (D) Decoding results for oboe F0 identification using rate.
